# A biosensor encompassing fusarinine C-magnetic nanoparticles and aptamer-red/green carbon dots for dual-channel fluorescent and RGB discrimination of *Campylobacter* and *Aliarcobacter*

**DOI:** 10.1101/2023.02.22.529591

**Authors:** Weixing Liu, Zhe Chi

## Abstract

The diarrhea pathogens *Campylobacter* and *Aliarcobacter* are similar in morphology and their resulting symptoms, making them difficult to be differentially diagnosed. Herein, we report a biosensor with two newly-synthesized modules to differentiate the genera-representative species of *C. jejuni* and *A. butzleri*. Module 1 was fusarinine C-decorated magnetic nanoparticles; module 2 consisted of *C. jejuni*-specific aptamer modified with red-emitting carbon dots (CDs) and *A. butzleri*-specific aptamer-modified green-emitting CDs. These two CDs had non-interfering spectra. Module 1 was used to selectively capture *C. jejuni* and *A. butzleri* from an un-cultured sample, and the specific CDs in module 2 would then recognize and bind to their counterpart bacteria when subjected to the collected module 1-bacteria complex. By measuring the fluorescence intensities from each CDs, the existence and abundance of each bacterium could be differentially indicated. This biosensor exhibited a wide detection range of up to 1 × 10^7^ CFU/mL and the lowest limit of detection (LOD) of 1 CFU/mL, for each bacterium. Thus, the biosensor with dual-fluorescent channels facilitated a culture-independent, ultrasensitive and discriminative detection of *C. jejuni* and *A. butzleri*. Remarkably, this fluorescent detection could be transformed into RGB color indication to render the visual discrimination. After integrating the microfluidics, this biosensor offered RGB differentiation of the two bacteria in human stool or chicken broilers with a LOD of 5 CFU/mL and turnaround time of 65 min. This work suggested a new biosensor-based methodology for the discrimination of *Campylobacter* and *Aliarcobacter* in real samples.

## 1. Introduction

Currently, the genera of *Campylobacter* and *Aliarcobacter* (formerly named as *Arcobacter*) [1], have been certificated as causes of gastrointestinal infections [2]. The infection of *Campylobacter*, mostly by *C. jejuni* and *C. coli*, causes campylobacteriosis that mainly manifests symptoms of bloody diarrhea, nausea, fever and abdominal cramps in humans [3, 4]; Whereas, the infection of *Aliarcobacter* induces a chronic watery diarrhea instead of the bloody one [5]. Moreover, campylobacteriosis has been implicated to associate with Guillain-Barré syndrome, a rare neurological disorder [6], colorectal cancer and inflammatory bowel disease [7]; *Aliarcobacter* infection has been associated with extraintestinal diseases like bacteremia [8]. Now, both *Campylobacter* and *Aliarcobacter* have been considered as emergent food- and water-borne pathogens and zoonotic agent responsible for bacterial diarrhoeal diseases [3, 8].

Notably, *Aliarcobacter* is close to *Campylobacter* with high similarity in structure and morphology [9]. Before the nomenclature as a new genus, *Aliarcobacter* was usually misidentified as atypical *Campylobacter* via conventional cultivation-dependent methods and phenotypic tests without sufficient sensitivity [10]. This could lead to underestimation of *Aliarcobacter* species in environmental samples and human [10] and in turn misdiagnosis, improper prescription, and delayed cure for the patients. Additionally, epidemiological data on *Aliarcobacter* are quite limited, and the incidence rate of the pathogen has been greatly underestimated [11]. More recently, rare cases of mixed etiology of *Campylobacter* and *Aliarcobacter* have been described [12, 13]. Therefore, approaches that allow discriminative detection between *Campylobacter* and *Aliarcobacter* are required. To date, several methods have been reported to distinguish *Aliarcobacter* from *Campylobacter* including conventional PCR [10], fluorescent in situ hybridization (FISH) techniques [10], DNA oligonucleotide arrays [14], and matrix-assisted laser desorption ionization–time of flight mass spectrometry (MALDI-TOF MS) [15]. However, PCR-based methods mostly require long-term bacteria isolation and pure culture as well as several hours’ of PCR reaction making them time-consuming and labor-intensive; the use of FISH, MALDI-TOF MS, and Raman spectroscopy suffer from consumption of expensive materials and the need for skilled technical staff, thus hindering them from general applications in food tests and clinical settings [11, 16]. Therefore, a novel sensing methodology for an accurate discrimination of *Campylobacter* and *Aliarcobacter* is still in demand to facilitate correct diagnosis and customized treatment for these two pathogens while assisting in their epidemiological surveillance worldwide.

Optical biosensors are effective and versatile analytical devices that can well perform highly selective detection towards biological targets including bacteria [17]. Optical biosensors have led to remarkable advances in clinical diagnosis, food control, and environmental monitoring [18]. Biosensors typically contain two basic functional units: a bioreceptor (e.g., enzyme, antibody or DNA) for selective recognition of the targets as well as a physico-chemical transducer (e.g., electrochemical, optical or mechanical) for translating this biorecognition into a signal [19]. This signal can be further read and recorded with a processing apparatus [18]. To be noted, within those optical sensing methodologies, fluorescence is particularly promising due to its high sensitivity, facile manipulation and ability to indicate analytes with low abundance [20]. Dual-channel fluorescent detection [21, 22] has value for the simultaneous sensing of two bacteria. This can also be achieved by employing two different fluorescent transducers with specific recognition to *Campylobacter* and *Aliarcobacter*, respectively. This strategy integrates sensing and transducing modules together to simplify the optical biosensors [23]. Moreover, these specific transducers can have minimal overlap in the emission spectra (e.g., green and red fluorophores) to avoid mutual interference. Upon binding to their respective bacterium, a specific fluorescence intensity from the transducer would correlate to the bacterial cell density, and this fluorescence signal can be represented by RGB color to help visualize the existence and abundance of the bacterium [24]. The RGB color scale can report the co-existence of *Campylobacter* and *Aliarcobacter*. To reduce this idea to practice, aptamer-conjugated carbon dots (CDs) [25-27] have value as fluorescent transducers for pathogenic bacteria detection: They combine the high specificity of aptamers [28] with the good photoluminescent properties of CDs [23].

It should be reiterated that real samples contain complicated matrices [29], which could lead to fluorescence interference [30]. Bacteria can be separated from the interfering matrices via magnetic nanoparticles (MNPs) decorated with aptamers [30] or antibodies [29]. Thus, specific and simultaneously capture of *Campylobacter* and *Aliarcobacter* is expected. To render a dual-bacteria selectivity to MNPs, a ligand that possesses affinity to both of the bacteria can serve as a functional molecule for chemical modifications. *Campylobacter* has an active transport system to uptake a xenosiderophore of enterobactin (Ent) in its iron-containing form despite its inability to synthesize siderophores [31]. Moreover, homologous genes of this system, e.g., *cfrA* and *ceuD* [31], can also be discovered via the mining of published *Aliarcobacter* whole genome sequences [32], thus, implying that this bacteria genus can also actively transport Ent. These suggest that Ent may serve as an active targeting molecule towards both *Campylobacter* and *Aliarcobacter*. Hence, the surface modification on MNPs with Ent is very likely to produce a tool for selective capture of these two bacteria. However, Ent has no active groups for easy chemical modifications. The fusarinine C siderophore (FsC) has active amine groups to offer sites for facile chemical reactions [33], which could simplify the synthesis of siderophore derivatized MNPs. There is currently no investigation into the active targeting property of FsC towards *Campylobacter* and *Aliarcobacter* making this issue interesting to explore.

Based on these considerations, a fluorescent biosensing system is designed to detect and discriminate *Campylobacter* and *Aliarcobacter* simultaneously in a dual-channel mode (Fig. 1). Briefly, *C. jejuni* and *A. butzleri* are taken as the representative species for *Campylobacter* and *Aliarcobacter*. FsC[Fe^3+^] chelates are used to modify MNPs, aiming to obtain the capture module 1 selective for both *C. jejuni* and *A. butzleri*. Meanwhile, red-emitting CDs are modified with the *C. jejuni-* specific aptamer and green-emitting CDs are modified with the *A. butzleri*-specific aptamer, in an attempt to prepare the bifunctional recognizing and transducing module 2. The two CDs within, would specifically bind to their counterpart bacterial cells captured by module 1, and correspondingly indicate the bacterial intensities with different intensities of red or green fluorescence to establish their correlations. Further, the fluorescence readouts are planned to be transformed into RGB values of red or green color schemes. At the different abundance of *C. butzleri*, the addition of their corresponding RGB values would produce a color grid as a reference. For a sample containing *C. jejuni* and/or *A. butzleri*, a specific color is expected to be produced after it is subjected the biosensor. By visual comparison of this color to the reference, it would be easy to estimate the abundance of the two bacteria in this sample, and obtain the specific bacterial densities by calculating the fluorescence intensities corresponding to this RGB value. In this work, the effectiveness for this designed biosensing approach to differentiate *C. jejuni* and *A. butzleri* from pure-culture and real samples, would be specified in detail.

**Fig. 1.**
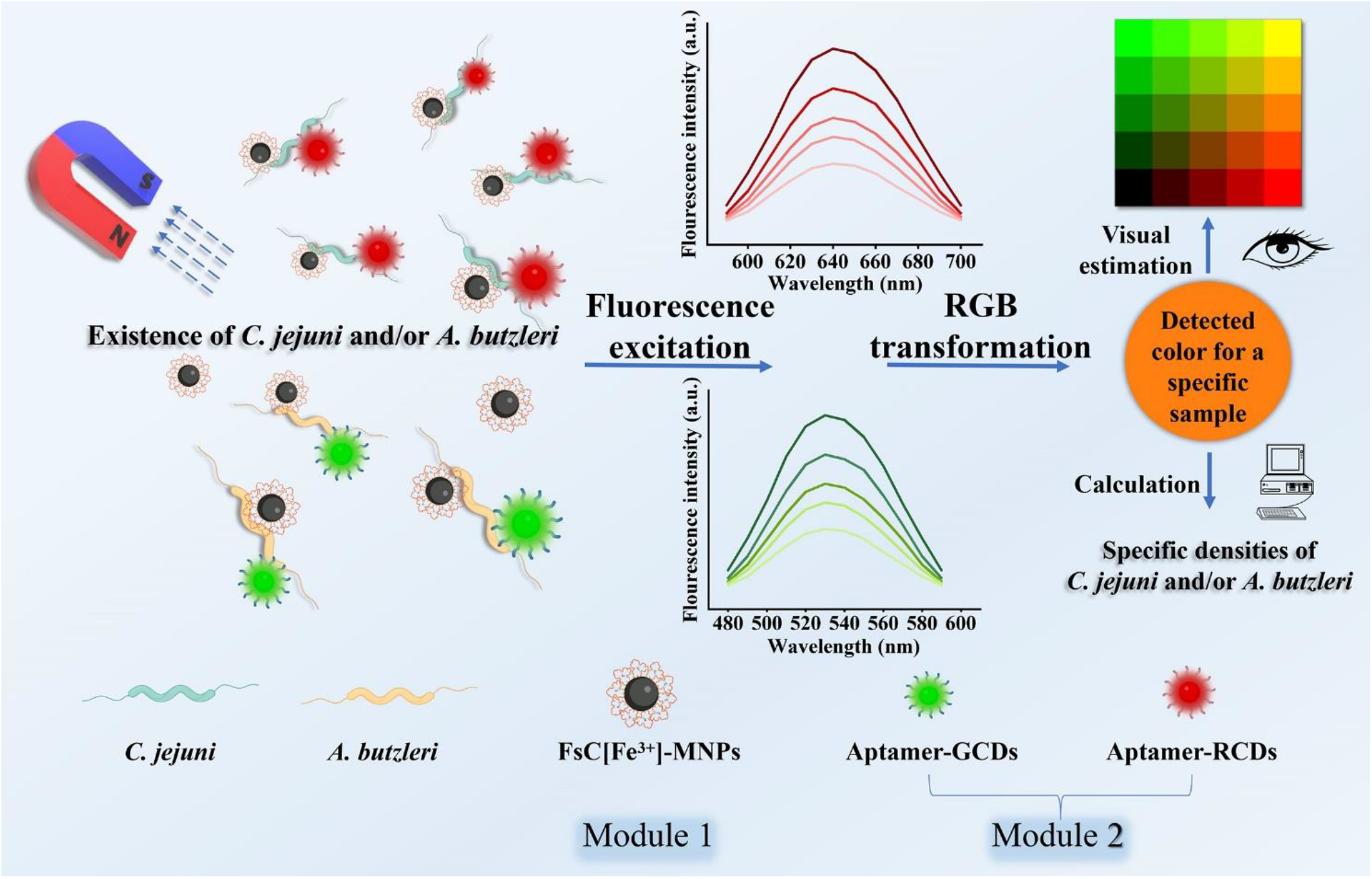
Schematic illustration for the designed modules and hypothetical working principle of the biosensor designed in this work.

## 2. Materials and Methods

The materials, the methods for synthesizing and characterizing each module of the biosensor, and the procedures for the absolute quantification of *C. jejuni* and *A. butzleri* are presented in the Supplementary Materials.

### 2.1. Selection and truncation of *A. butzleri*-specific aptamer

#### 2.1.1 Selection of *A. butzleri*-specific aptamer

Selection of the *A. butzleri*-specific aptamer was performed using high-throughput sequencing (HTS)-assisted SELEX [30] with the oligonucleotide ssDNA library comprising a central random region of 40 nucleotides (nt) flanked on both sides by 20 nt primer regions (sequence: 5’-TGGACCTTGCGATTGACAGC (40 nt) GCAGACATGAGTCTCAGGAC-3’). For the counter selection, a bacterial community encompassing *C. jejuni, H. pylori, C. coli, L. casei, B. bifidum, L. johnsonii, E. faecalis, E. coli, S. aureus, S. enteritidis, L. monocytogenes, S. gordonii, R. pickettii*, and *S. dysenteriae* was used at the first, third, sixth, ninth, and twelfth rounds. The ssDNA library from the twelfth round was sequenced via HTS performed by Novogene Co. Ltd. (Beijing, China); the top-five abundant sequences given by the HTS were tested for affinity and specificity against the bacterial community to recognize the most suitable aptamer candidate referring to the former work [30].

#### 2.1.2 Truncation of *C. jejuni*-specific and *A. butzleri*-specific aptamers

The sequence of *C. jejuni*-specific aptamer was described in our previous study [34]. The truncation of *C. jejuni*-specific and *A. butzleri*-specific aptamers was performed as described previously [35, 36]. Briefly, DNAMAN was used to predict secondary structure of these aptamers; the predicted binding motif regions from the original aptamers were truncated [36] and individually synthesized. The optimal truncated aptamer was determined by their affinities and specificities for *C. jejuni* and *A. butzleri* versus those for the bacteria in the community.

### 2.2. Targeting property of FsC[Fe^3+^] to *C. jejuni* and *A. butzleri*

FsC[Fe^3+^] was labelled by 6-carboxyfluorescein (6-FAM) (Thermo Fisher Scientific, Waltham, USA) to indicate the targeting of FsC[Fe^3+^] to *C. jejuni* and *A. butzleri*. To synthesize it, FsC[Fe^3+^] was conjugated to the free carboxyl group of the 6-carboxyfluorescein (6-FAM) via its amine groups at a molar ratio of 1: 2. In detail, 1.5 mg 6-FAM was reacted with 1.0 mg FsC[Fe^3+^] under the catalysis of 1.9 mg EDC·HCl and 2.3 mg NHS in 1.0 mL of 15 mM MES buffer (pH=6) at 25°C for 2 h. Excess dye was removed via dialysis (MW cut-off: 500–1000) at 4°C for 24 h, and FsC[Fe^3+^]-FAM conjugates were solidified by freeze drying. The fluorescence intensity of FsC[Fe^3+^]-FAM and FsC[Fe^3+^] (excitation wavelength 492 nm, emission wavelength 514 nm) were immediately detected using a fluorescence microplate reader (TECAN Infinite® M200 PRO, Switzerland). The targeting of FsC[Fe^3+^]-FAM to *C. jejuni* and *A. butzleri* started with a cell density of 1.0 × 10^7^ CFU/mL *C. jejuni, A. butzleri*, and *E. coli* incubated with 1 mg of FsC[Fe^3+^]-FAM. The sample was then observed under a confocal laser microscope with an excitation wavelength of 492.0 nm and an emission wavelength of 514.0 nm. Next, 1 mg of FsC[Fe^3+^]-FAM was incubated with *C. jejuni, A. butzleri*, and the control bacteria of *E. coli, R. pickettii, S. enteritidis, L. johnsonii, E. faecalis* and *B. bifidum* (each at the density of 1.0 × 10^7^ CFU/mL) at 37°C for 15 min. The mixtures were centrifugated, washed three times with 1× PBS buffer, and then analyzed with a fluorescence microplate reader (TECAN Infinite® M200 PRO, Switzerland) (excitation wavelength: 492.0 nm, emission wavelength: 514.0 nm).

### 2.3. Optimization of working conditions for the biosensor

The working conditions of module 1 and 2 were optimized, including the concentrations of module 1 and module 2 to react with *C. jejuni* and *A. butzleri*, as well as the time required for module 1 to capture *C. jejuni* and *A. butzleri*, and module 2 to indicate the bacteria. In detail, module 1 with a series of final concentrations (0, 100, 200, 300, 400, 500, 600 and 700 µg/mL) were incubated with 1.0 × 10^8^ CFU/mL of *C. jejuni* or *A. butzleri* in 1.0 mL 1 × PBS buffer at 37°C for 60.0 min, respectively. Magnetic separation was performed next, and the number of bacteria captured by module 1 was measured by absolute quantification using qPCR.

The incubation time was optimized next by incubating module 1 at the optimal concentration with 1 × 10^8^ CFU/mL of *C. jejuni* or *A. butzleri* in 1.0 mL 1 × PBS buffer for 0, 10, 20, 30, 40, 50 and 60 min. The number of captured bacteria was determined by measuring the *C. jejuni* and *A. butzleri* left in the supernatants via qPCR. To optimize the working conditions for module 2, the optimal concentrations of each CD were determined by incubating ARCDs or AGCDs at a series of final concentrations with their counterpart bacteria (1×10^7^ CFU/mL) at 37°C for 20 min in 1.0 mL 1 × PBS buffer. The amount of ARCDs was 2.5, 5, 7.5, 10, 12.5 and 15 µg and that for AGCDs was 20, 40, 60, 80, 100 and 120 µg, respectively. The bacteria were collected by centrifugation, washed twice, and analyzed with a fluorescence microplate reader (TECAN Infinite® M200 PRO, Switzerland). Using the optimal concentrations determined above, each CD was reacted with its specific bacteria for various times (0, 10, 20, 30, 40, 50 and 60 min) and tested for fluorescence as described above to obtain the optimal reaction time.

### 2.4. Detection of *C. jejuni* and *A. butzleri* in real samples

The microfluidics-assisted detection of *C. jejuni* and *A. butzleri* in real samples used the methods we published previously [30]. The preparation of real samples can be seen in the supplementary materials.

### 2.5. Absolute quantification of *C. jejuni* and *A. butzleri* by qPCR

The absolute quantification of *C. jejuni* was determined according to the copy number of the *C. jejuni gyrA* gene [37] via qPCR [34], yielding a standard calibration curve of [y =−2.3308x +36.303, R² = 0.9919] where x is the log value of the plasmid copy number, and y is the corresponding C_T_ value. *A. butzleri* was similarly quantified by qPCR according to the copy number of the *A. butzleri hsp60* gene [38], thus yielding a stand curve of [y = −2.9003x + 38.808, R² = 0.9985].

### 2.6. Statistical analysis

Experiments were conducted in triplicate (n = 3) or sextuplicate (n = 6) as required. The results were expressed as mean ± standard deviation. Comparative studies of means were performed using one-way ANOVA; p < 0.05 was considered statistically significant.

## 3. Results and Discussion

### 3.1. FsC[Fe^3+^]-modified Fe_3_O_4_ MNPs as the capture module

Previously, MNPs decorated with specific ligands, e.g., aptamers [39] and antibodies [16], have been used to capture and separate *C. jejuni* from matrices *in vitro*. However, we intended to capture *C. jejuni* and *A. butzleri* simultaneously by using the MNPs modified with a ligand specific for both of the two bacteria (**Fig. 1**). As proposed above, FsC might be able to serve as this specific ligand. To verify this hypothesis, the targeting ability of FsC to *C. jejuni* and *A. butzleri* was testified with a fluorescent probe of 5-carboxyfluorescein-FsC conjugate (**Fig. S1a**). As shown in **Fig. 2**, the FsC[Fe^3+^]-FAM probe could label both *C. jejuni* and *A. butzleri* with bright green fluorescence but not *E. coli* controls. This result clearly justifies that FsC[Fe^3+^] can be actively imported by *C. jejuni* and *A. butzleri*, but not by *E. coli*, indicating that FsC targets *C. jejuni* and *A. butzleri* via the siderophore transport systems of these two bacteria based on xenosiderophore uptake. Further, it was verified that other commonly-isolated intestinal bacteria could not actively import FsC[Fe^3+^]-FAM as manifested via the extremely low fluorescence emitted from the bacterial cells after incubation with FsC[Fe^3+^]-FAM probe (**Fig. S1b**).Overall, it can be validated that FsC[Fe^3+^] can be actively transported by both *C. jejuni* and *A. butzleri* as a targeting molecule, suggesting that FsC[Fe^3+^] can be used as a ligand to modify MNPs to synthesize a selective capture module for these two bacteria.

**Fig. 2.**
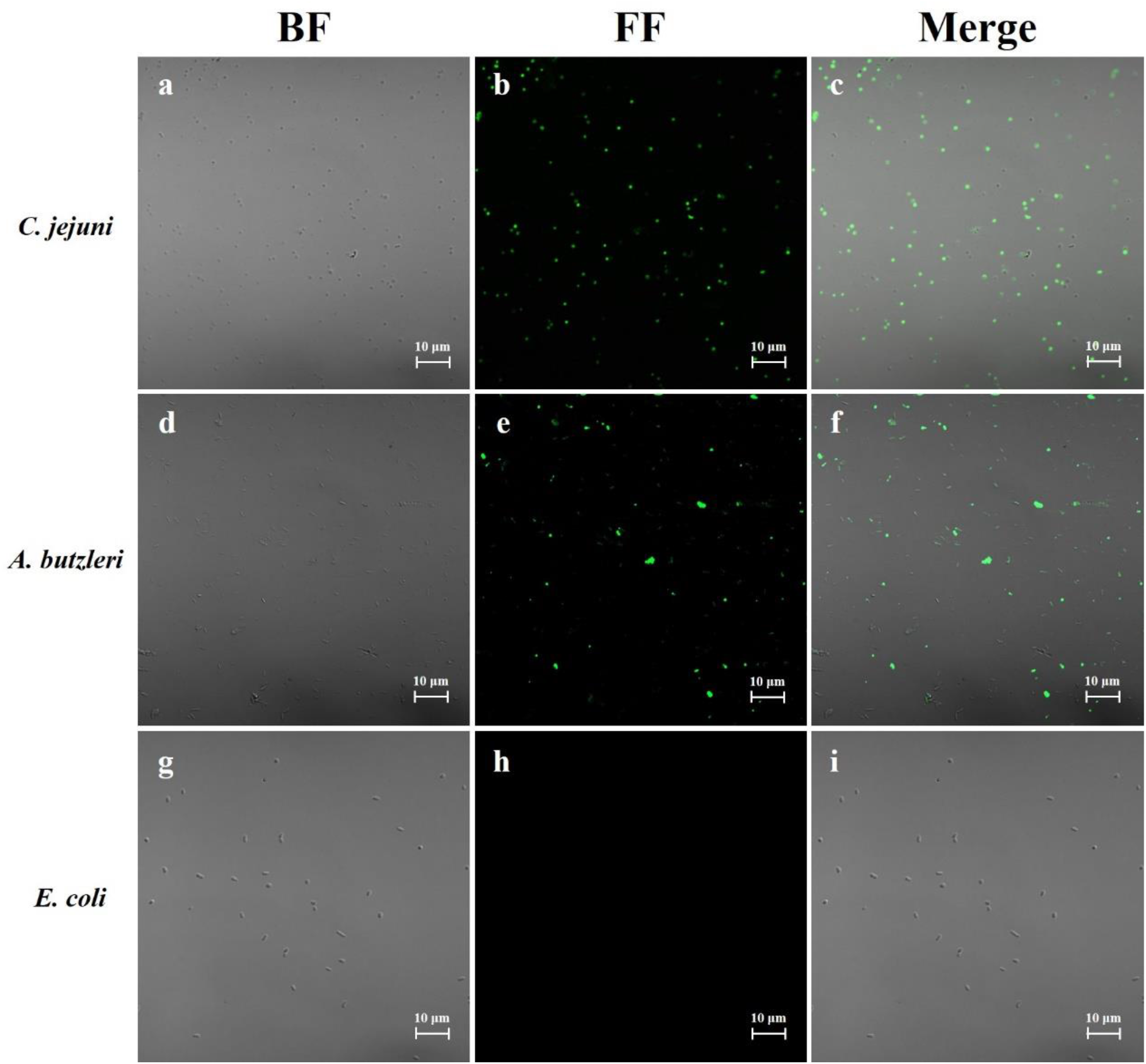
Targeting ability of FsC[Fe^3+^]-FAM conjugates to the tested bacteria by the observation of confocal laser microscopy. (a, b, c) Images for FsC[Fe^3+^]-FAM incubated with *C. jejuni* under bright field (BF), fluorescence field (FF), and their merged image. (d, e, f) Images for *A. butzleri* after incubation with FsC[Fe^3+^]-FAM under BF, FF, and the merged image. (g, h, i) Images for the labelling of FsC[Fe^3+^]-FAM to *E. coli* under the same fields. Scale bar: 10.0 µm. Excitation wavelength (λex) = 492.0 nm, emission wavelength (λem) = 518.0 nm.

Previously it was clarified that *C. jejuni* can actively import the iron-chelating enterobactin (Ent[Fe^3+^]) siderophore [40, 41], however, it is unclear whether *A. butzleri* can uptake any siderophore. However, the enterobactin commodity is costly, and the only chemically reactive site on the enterobactin ring is the phenolic hydroxyl group, which would make chemical modifications for this moiety suffer from much difficulties. Nevertheless, it was shown here for the first time that the fungal FsC siderophore can also be actively transported by *C. jejuni* and *A. butzleri*. More remarkably, FsC has amine groups that are more active and facile and can be chemically modified. FsC is more easily available and inexpensive via published preparation protocols [33]. Thus, it would be realistic to use FsC[Fe^3+^] to modify MNPs to produce a commercial product that captures bacteria. The appearance would change from the colorless empty FsC to dark red of FsC[Fe^3+^] upon chelating Fe^3+^ [33]. This feature, together with the bacteria-targeting of FsC, may help it develop into new recognizing or transducing modules that develop other novel biosensors for detecting bacteria.

To obtain an effective capture module of FsC[Fe^3+^]-MNPs, MNPs with an average particle size of 6.67 ± 3.57 nm were initially prepared (**Fig. 3a**), following our previous work [30]. The MNPs were then modified by FsC[Fe^3+^]. The FT-IR analysis (**Fig. 3b**) of the magnetically collected product showed the appearance of a new characteristic peak at 1395.6 cm^−1^ in its spectrum (red), which was ascribed to a N–H bond vibration [42]; while, the same peak was not discovered in the spectrum of plain MNPs (blue). These results indicated that FsC[Fe^3+^] was successfully conjugated with MNPs via amido bonds. However, when these FsC[Fe^3+^]-MNPs were incubated with a bacterial community constructed artificially (see experimental section in Supplementary Materials). Their selectivity to *C. jejuni* and *A. butzleri* were not obvious, and many other bacteria can be captured by the FsC[Fe^3+^]-MNPs.

**Fig. 3.**
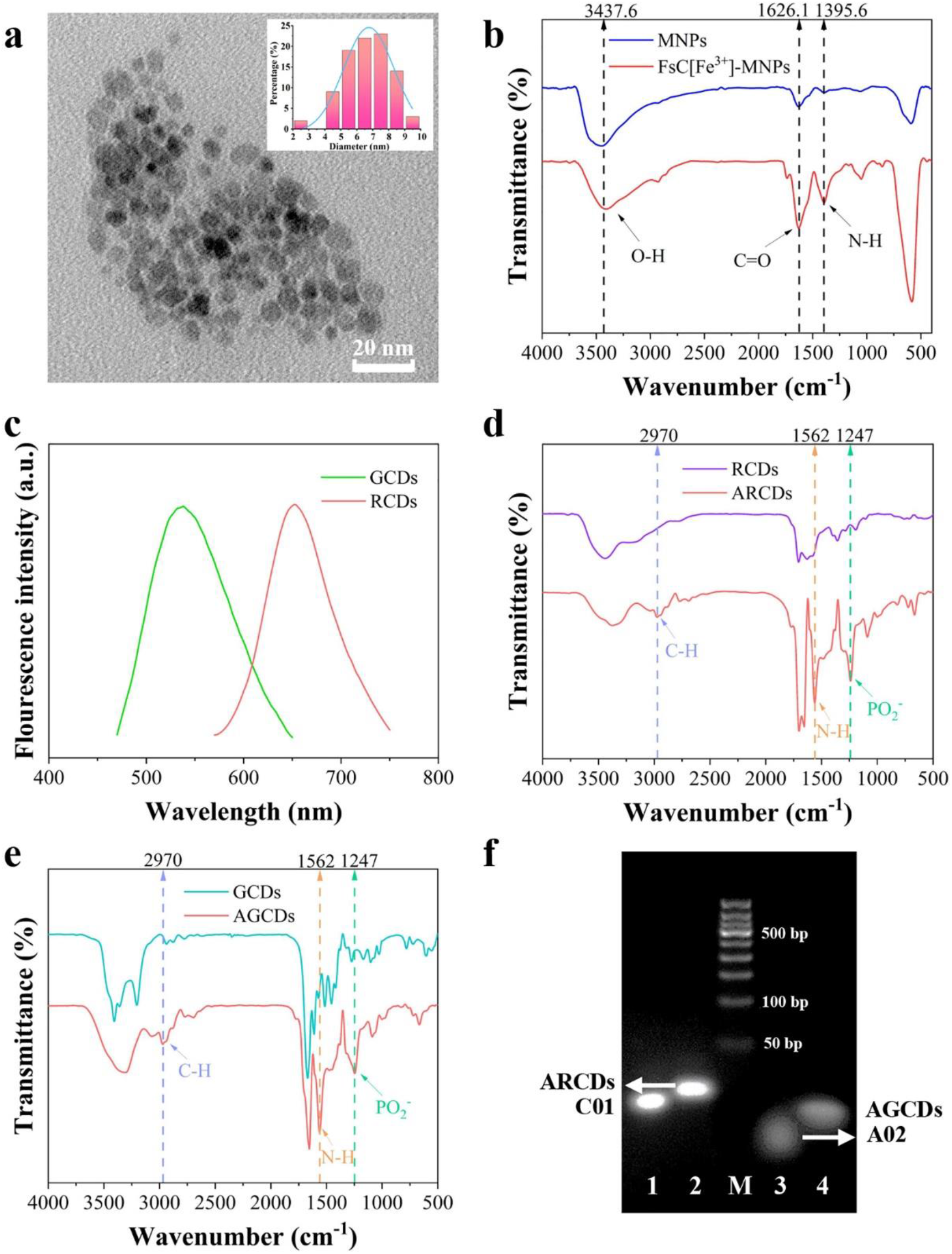
(a) TEM observation and the average diameter (inset) of synthesized magnetic nanoparticles (MNPs). (b) FT–IR spectra of the original (blue) and FsC[Fe^3+^]-modified MNPs (red). (c) Fluorescence emission spectra of RCDs and GCDs. (d-e) FT-IR spectra of the original red-emitting CDs (RCDs) and green-emitting CDs (GCDs) as well as *C. jejuni*-specific aptamer-modified RCDs (ARCDs) and *A. butzleri*-specific aptamer-modified GCDs (AGCDs). (f) The electrophoretic mobility shift for the original *C. jejuni*-specific aptamer C01 (Lane 1), C01-modified ARCDs (Lane 2), the original *A. butzleri*-specific aptamer A02 (Lane 3), and A02-modified AGCDs (Lane 4); Lane M: DNA ladder.

Z-potential measurements of FsC[Fe^3+^]-MNPs showed that they were negative (−11.2 mV) due to the unreacted free carboxyl groups on FsC[Fe^3+^]-MNPs. This charge led to low selective adhesion of FsC[Fe^3+^]-MNPs to bacteria via electrostatic interactions [30]. To alleviate this non-specific binding, CaCl_2_-blocking [43] was used to neutralize the residual surface charges of FsC[Fe^3+^]-MNPs. After FsC[Fe^3+^]-MNPs were treated with 5.0% (w/v) CaCl_2_ (Z-potential increased to 2.3 mV), they could have higher selectivity for *C. jejuni* and *A. butzleri* in the same bacterial community as above. The capture efficiency was 45.0% for *C. jejuni*, 42.2% for *A. butzleri*, and less than 13.0% for the other bacteria in total (**Fig. S2a**). Thus, Ca^2+^-doped FsC[Fe^3+^]-MNPs (Ca/FsC[Fe^3+^]-MNPs) are an effective tool for the simultaneous and selective capture of *C. jejuni* and *A. butzleri*; these were verified to have value in module 1. The optimal concentration and time needed for module 1 to fully capture *C. jejuni* and *A. butzleri* were 500 µg/mL and 30 min, respectively (**Fig. S2b and c**).

### 3.2. Aptamer-modified CDs as the bifunctional recognizing and transducing module

As designed, *C. jejuni*-specific aptamer modified red-emitting carbon dots (ARCDs) and *A. butzleri*-specific aptamer modified green emitting AGCDs, were united as a bifunctional recognizing and transducing module to differentiate *C. jejuni* and *A. butzleri*. To achieve it, red CDs (RCDs) and green CDs (GCDs) were correctly synthesized (**Fig. S3**). It could be found that their fluorescence spectra only slightly overlapped from approximately 570 nm to 620 nm, and the distance for their maximal emission wavelength (640.0 nm of RCDs vs 530.0 nm of GCDs) was long (**Fig. 3c**), which indicated that the two CDs did not interfere with each other in fluorescence emission. This confirms the value of using their fluorescence as separate signal channels.

As for the aptamers, the sequence of the *C. jejuni*-specific aptamer was referred to our former work [34], and the selection and identification of the *A. butzleri*-specific aptamer were completed here **(Fig. S4**). Both of the aptamers were truncated, and the affinities of the resulting shortened aptamer candidates to their corresponding bacteria were determined to obtain the most specific one to *C. jejuni* or *A. butzleri* (**Fig. S5 and S6**). The modified CDs were collected and characterized after the CDs reacted with their aptamer counterparts. The fluorescence excitation and emission spectra of the aptamer-modified ARCDs and AGCDs showed no change at first. The FT-IR analysis (**Fig. 3d and e**) indicated new characteristic peaks at 2970 cm^−1^ and 1247 cm^−1^, which were ascribed to C–H [42] and PO_2-_ [44] bond vibrations in the spectra of modified CDs, respectively; these peaks were not found in the spectra of original RCDs and GCDs. The EMSA assay (**Fig. 3f**) for ARCDs and AGCDs showed less migration of ARCDs and AGCDs in electrophoresis versus those in aptamers. These assays collectively justified that the bacteria-specific aptamers were conjugated to the respective CDs. And, the specificity of the optimized ARCDs and AGCDs to their counterpart bacteria was also confirmed (**Fig. S7**). Moreover, the maximal grafting ratios for aptamers to ARCDs and AGCDs were testified to be 18.8 pmol/µg and 2.6 pmol/µg (**Fig. S8a and b**), respectively.

The efficiency of ARCDs and AGCDs for indicating the densities of their targeted bacteria were investigated next. Here, each CD was incubated with its bacterium counterpart separately in an ideal environment of 1 mL 1 × PBS buffer. The optimal amount of ARCDs was 5 µg when using 1 × 10^7^ cells of *C. jejuni* (**Fig. S8c**); the optimal reaction time was 10 min (**Fig. S8d**). Further, after ARCDs were incubated with *C. jejuni* increased from 10 to 1 × 10^7^ CFU/mL, the determined intensity of red fluorescence grew accordingly (**Fig. 4a**). Importantly, interfering fluorescence might be emitted from *C. jejuni* under the excitation wavelength of ARCDs as background While, in fact, it was revealed that the use of red-emitting ARCDs did prevent this interference after determination. Thus, it could be plotted that the logarithms of the fluorescence intensities and cell densities exhibited a linear correlation, conforming to a linear regression equation of [y = 0.2684x +1.5452, R^2^ = 0.9927], as shown in **Fig. 4b**. This demonstrates that ARCDs can bind to the *C. jejuni* cells with high affinity, thus indicating a range of 10 to 1 × 10^7^ CFU/mL.

**Fig. 4.**
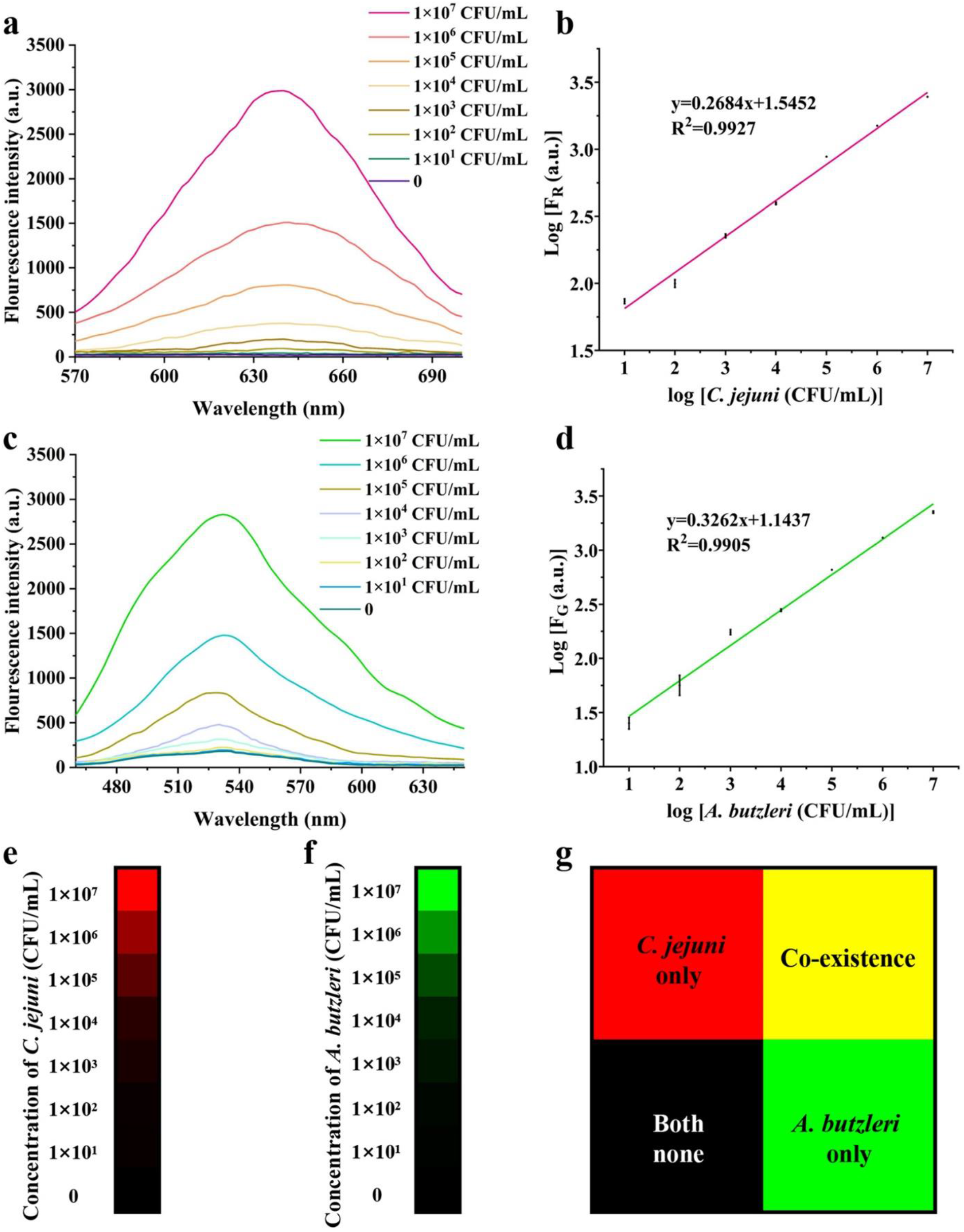
(a, b) Detected fluorescence spectra for ARCDs to indicate *C. jejuni* at 0, 10, 10^2^, 10^3^, 10^4^, 10^5^, 10^6^, and 10^7^ CFU/mL as well as their linear correlation. (c, d) Detected fluorescence spectra for AGCDs to indicate *A. butzleri* at 0, 10, 10^2^, 10^3^, 10^4^, 10^5^, 10^6^, and 10^7^ CFU/mL as well as their linear correlation. Data, where necessary, are presented as mean ± SD; n = 3. (e-g) Mosaic bar generated by transforming detected fluorescence intensities into RGB colors against different cell densities of *C. jejuni* and/or *A. butzleri*.

Likewise, *A. butzleri* could be indicated by AGCDs (the optimal amount of AGCDs was 40 µg for 1 × 10^7^ cells in 1 mL PBS; the optimal reaction time was 10 min as shown in **Fig. S8e and f**. Moreover, there was negligible bacterial autofluorescence at the emission wavelength of AGCDs. Thus, an indicating range of 10 to 1 × 10^7^ CFU/mL could be achieved, conforming to an equation of [y = 0.3262x +1.1437, R^2^ = 0.9905] (**Fig. 4c and d**). With the equations, it can be calculated that the theoretical values of limit of detection (LOD) are both 1 CFU/mL for *C. jejuni* and *A. butzleri* when module 2 is used as the transducer. The LOD here is the lowest as compared with those reported in previous work regarding the culture-independent discriminative detection between *Campylobacter* and *Aliarcobacter*, which may include those of 10^2^ for *C. jejuni* and 10^2.5^ for *A. butzleri* from stool samples by using Real-Time multiplex PCR [38]; 10^2^ for *Campylobacter* and 10^3^ for *Aliarcobacter* from river water samples by using FISH [10]; and 10^4^ for *C. jejuni* from chicken samples by using DNA oligonucleotide arrays [14]. This progress reflects the ultra-sensitivity of module 2, owing to the high luminance of the CDs and their non-interfering emission spectra. Moreover, the cost for synthesized truncated aptamers and CDs are low, thus suggesting the possibility of using module 2 in a commercial product.

On obtaining the indication ranges, the fluorescence intensities were then transformed to colors as proposed. To do so, at the *C. jejuni* density of 1 × 10^7^ CFU/mL, the resulting fluorescence intensity was set as the maximum (100%), and it was set as the standard red color with the RGB value of 255, 0, 0. As indicated above, when the cell density at the rest 7 points descended, the corresponding fluorescence intensity would decrease accordingly. For each density point, the percentage of its tested fluorescence intensity accounting for the maximum could be calculated. With this percentage to time the red RGB value, a reduced RGB value representing a new color was generated. Thus, for all the cell density points, a series of discrete colors descending from red (RGB value: 255, 0, 0) to dark red, and finally to black, could be visualized as a mosaic bar (**Fig. 4e**). Similarly, the fluorescence intensities for *A. butzleri* ranged from 1 × 10^7^ to 0 CFU/mL could be transformed to a mosaic color bar changed from green (RGB value: 0, 255, 0) to dark green, and finally to black (**Fig. 4f**). As for the co-existence of the two bacteria at certain cell densities between 0 and 1 × 10^7^ CFU/mL, the addition of RGB values transformed from the detected fluorescence intensities would produce a new color, following the below equation:

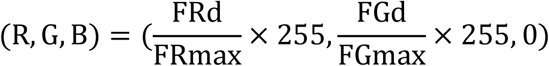

Here, FRd is the detected red fluorescence intensity, FGd is the detected green fluorescence intensity, FRmax is the maximal red fluorescence intensity after ARCDs reacted with 1×10^7^ CFU/mL *C. jejuni*, and FGmax is the maximal green fluorescence intensity after AGCDs reacted with 1×10^7^ CFU/mL *A. butzleri*.

For example, yellow color would be generated when the densities of the two bacteria were both 1 × 10^7^ CFU/mL; pure red or pure green indicates the existence of only *C. jejuni* or *A. butzleri*; black indicates that neither is present (**Fig. 4g**). Nevertheless, this is merely a theoretical visualization; the real performance of the RGB illustration for the co-existence of *C. jejuni* and *A. butzleri* should be clarified when the bacteria and ARCDs/AGCDs are mixed together in the same system and then captured by module 1 of Ca/FsC[Fe^3+^]-MNPs.

### 3.3. Working principle of the dual-channel biosensor

When the two bacteria and CDs reside in the same solution, possible non-specific binding for a CD to its non-targeted bacterium (e.g., ARCDs to *A. butzleri* and AGCDs to *C. jejuni*) may take place, which is usually caused by the size effect of CDs and the electrostatic interactions between CDs and bacteria [23]. In this context, at the exciting wavelength of one CD in this work, fluorescence may be emitted from the bacteria cells-non-specific CDs complexes, producing interference. To verify this concern, *C. jejuni* was incubated with AGCDs and *A. butzleri* was incubated with ARCDs respectively in separate solutions. Consequently, the suspicious fluorescence interference was confirmed to exist, and their specific intensities at every cell density of each bacterium was also determined (**Fig. 5a and b**). It could be found that the influence of the interfering fluorescence began to occur when the bacterial density was above 1 × 10^3^ CFU/mL. Importantly, module 1 was designed to capture the two bacteria from samples, and the module 1-bacteria complex was subjected to the binding by module 2. In this way, possible interfering noise came from module 1 could not be ignored when the fluorescence of module 1-bacteria-module 2 complex was measured. It was determined that such interfering fluorescence did exist when module 1 was excited under the wavelength of AGCDs, while it was neglectable when excited under the wavelength of ARCDs (**Fig. S9**). Thus, this specified noise should be subtracted to correct the signal readouts.

**Fig. 5.**
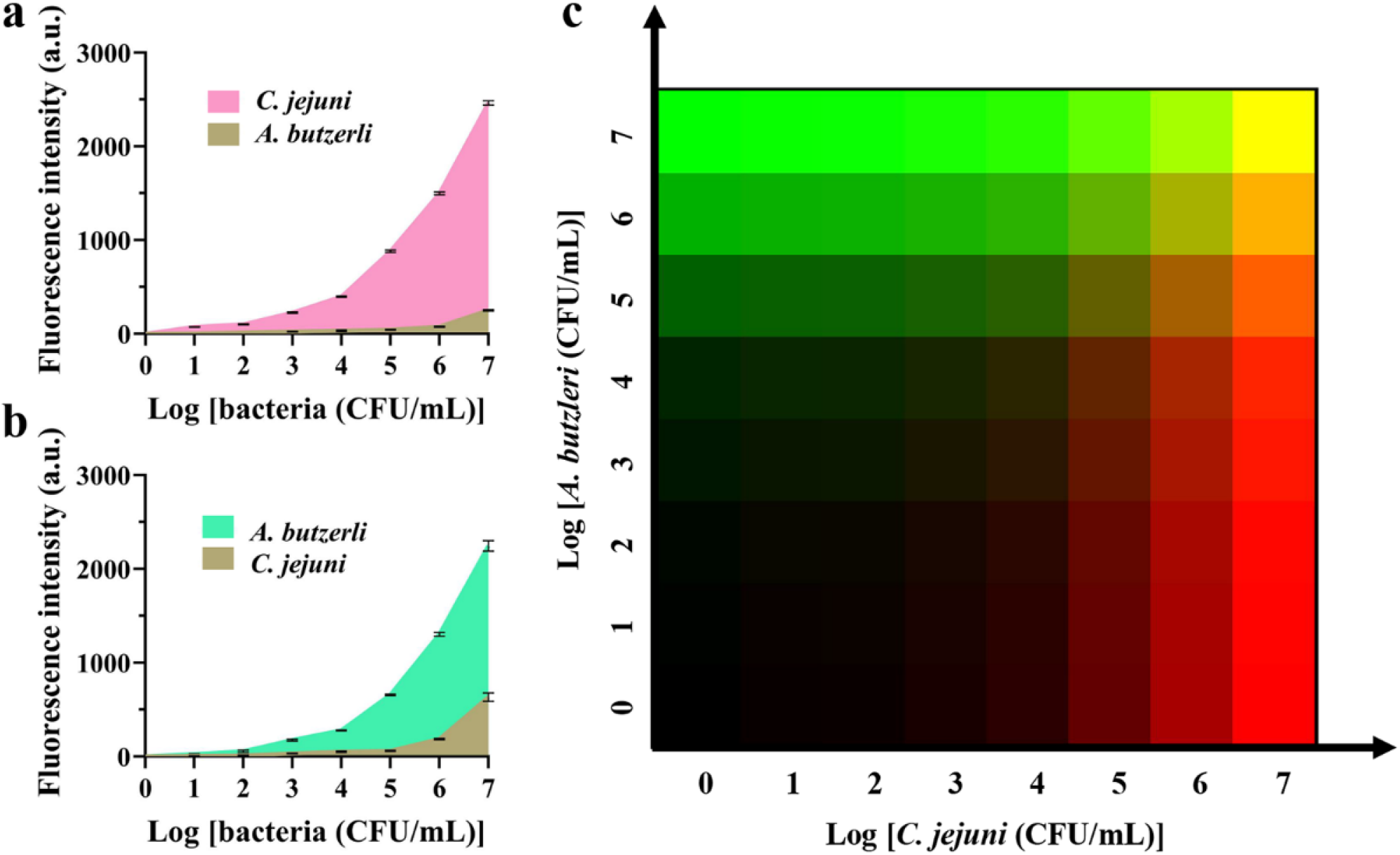
(a, b) Intensities of fluorescence background from non-specific binding of ARCDs to *A. butzleri* and AGCDs to *C. jejuni* (brown) merged in the detected florescence from specific CD-bound bacteria (b). Intensities of fluorescence from non-specific binding of AGCDs to *C. jejuni* (brown) merged in the detected florescence from specifically-bound ARCDs and *C. jejuni*. Data are presented as mean ± SD, n = 3. (c) The corrected 8 × 8 RGB color grid indicating the existence of *C. jejuni* and/or *A. butzleri* with different cell densities.

After the correction, a precise 8 × 8 RGB color grid (**Fig. 5c**) was plotted to indicate the existence and density difference of *C. jejuni* and *A. butzleri*. Both ranged from 0 to 1 × 10^7^ CFU/mL in the same sample after they were captured by magnetic separation and detected by a red/green dual-channel measurement. This confirmed the working principle of the biosensors. In brief, a sample suspected to contain *C. jejuni* and/or *A. butzleri* was treated with module 1 to enrich the bacteria. The module 1-bacteria complex was then collected by magnetic separation and transferred to interact with module 2. After incubation, the module 1-bacteria-module 2 complex was again collected by magnetic separation and subjected to a dual-channel fluorescent measurement. Afterwards, a new color was initially produced, and it can be referenced to the above color grid to judge the existence and estimate the approximate density of each bacterium in a more sensitive and rapid way by simple and direct visual observation. Meanwhile, corresponding to this visual estimation, the specific cell densities of *C. jejuni* and *A. butzleri* can be calculated with the obtained equations to validate this naked-eye recognition.

Unlike the case of using parallel fluorescent and colorimetric channels as a twin-modal sensing for one bacterium [45], the work here employs two fluorescent channels for sensitive detection of two bacteria. To note, a novel work has presented a multi-channel fluorescent biosensing platform for the discriminative identification of 10 bacteria simultaneously, with the lowest LOD of 2.5 × 10^5^ CFU/mL [46]. Despite the inadequate sensitivity, this work is inspiring because it can stratify between closely related bacteria. Herein, our work shows culture-independent, ultrasensitive and more importantly, discriminative detection of *C. jejuni* and *A. butzleri* by introducing a dual-channel fluorescent biosensor that can facilitate the transduction from the abundance difference of these two bacteria to fluorescence signals, as well as the further transformation to visualized RGB color indication and differentiation. To note, the RGB colors here is transformed from the fluorescent signals, and they cannot be read by a smartphone implanted a color-recognition software like ImageJ [45] However, in another way, the emitted fluorescence can be taken as an image by a smartphone; this image can be analyzed with a specialized software implanted in the smartphone and transformed in RGB presentation [27]. This approach would offer visual detection of multiple analytes, including different bacteria, with portable devices.

### 3.4. Microfluidics-assisted RGB discrimination of *Campylobacter* and *Aliarcobacter* in real samples

Further, the effectiveness of this biosensor for the application in real samples was evaluated. According to the previous studies [14, 34, 38], raw chicken and human stool were used as representative real samples to evaluate the effect of RGB discriminative detection for *C. jejuni* and *A. butzleri* with the dual-channel fluorescent biosensor in this work. Whereas, the matrices of chicken and stool may also cause strong fluorescence interference under the exciting wavelength of ARCDs and AGCDs. Hence, the centrifugal microfluidic plate used previously by us [30] was adopted here to pre-treat the bacteria-containing real samples, which could eliminate the possible fluorescence interference. Afterwards, the detection was performed by the developed biosensor. Table S1 and S2 show that approximately 50%–65% of the two bacteria could be collected from both samples after the microfluidic pre-treatment as verified by absolute qPCR quantification. The spike recovery experiments were validated by PCR and showed 70.0% to 100.0% detection with an actual LOD of 5 CFU/mL for both bacteria (Table S3 and S4). Further, the RGB grid could also be readily obtained to indicate the difference in abundance of *C. jejuni* and *A. butzleri* in real samples (Fig. 6). In Fig. 6a, it could be obviously identified the bacterial difference in the human stool sample with red, green or yellow colors, when *C. jejuni* and *A. butzleri* were both at the concentrations between 1.0 × 10^4^ and 1.0 × 10^7^ CFU/mL.

**Fig. 6.**
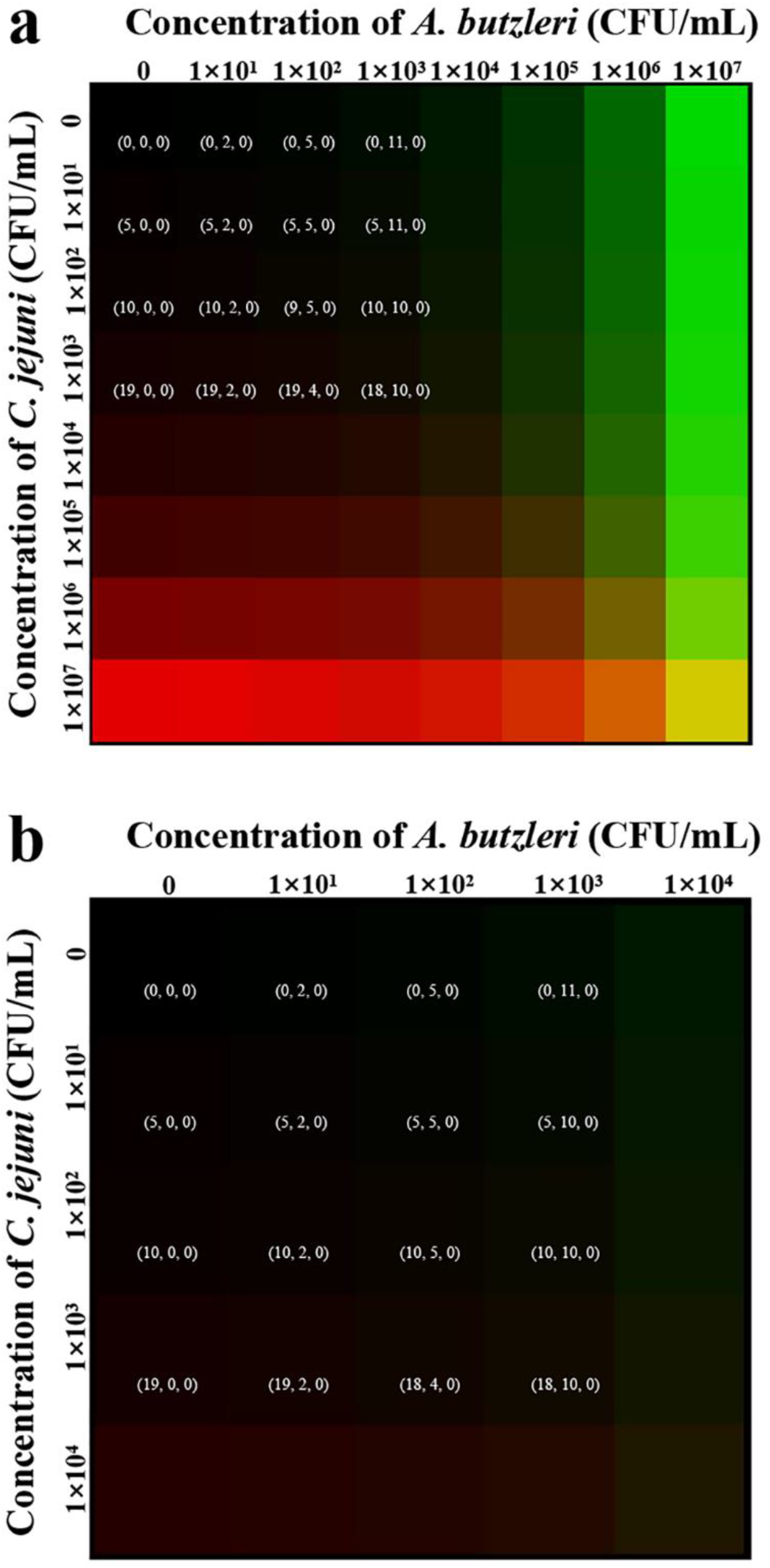
RGB color grid to indicate the existence of *C. jejuni* and/or *A. butzleri* in fecal samples (a) and chicken broiler samples (b). The RGB values for the close black-scheme colors, representing the bacteria with densities lower than less than 1 × 10^3^ CFU/mL, were specified on the color grid to show the difference.

However, when the cell density of these two bacteria were 0-1.0 × 10^3^ CFU/mL in human stool and chicken samples, the color identification was gradually covered by the black (Fig. 6a and 6b). Specially, bacteria above 1.0 × 10^3^ CFU/mL in chicken samples were not detected, due to maximal 1000 CFU/g carcass is allowed in frozen broiler as regulated by the European Commission [47], and that of *A. butzleri* was referred to it as no specific regulation for it has been published. In this situation, it was hard to distinguish these black colors to estimate the existence of each bacterium by simple naked-eye identification. Nevertheless, these black-scheme colors still hold significance for they can indicate that the real samples may not contain enough *C. jejuni* or *A. butzleri* that can cause illness or can be positively diagnosed as the pathogens for diarrheal patients. To enable the visual estimation for the bacteria in low abundance, the RGB values for these few black colors, were still specified in the grid as references. Thereby, a specific RGB value for a detected sample, which fall into the relatively black areas, can be compared with these referring values to evaluate the approximate densities of the two bacteria, thus to confirm the corresponding density after calculated using the equations above with the RGB value.

Herewith, it can be noted that the discrimination of *C. jejuni* and *A. butzleri* in both pure-culture and real samples using the RGB color reference is more sensitive and rapid than searching a datasheet recording large numerical arrays of fluorescence intensities (Table S3 and S4). Additionally, it takes approximately 20 min for the microfluidic pre-treatment and 40 min for the biosensing detection; plus, about 5 min is needed for the interval procedures, such as adding and pipetting samples, magnetic separation and operating equipment. Totally, it requires 65 min to finish one round of microfluidics-assisted dual-channel fluorescent biosensing discrimination of *C. jejuni* and *A. butzleri*, indicating it is a rapid detection platform. Besides, the capture capability of module 1, and specificities of above-identified aptamers for different species of *Campylobacter* and *Aliarcobacter* are without significant difference (**Fig. S10**), suggesting its genus-selectivity rather than perfect analyte-specificity [48]. This selectivity is significant because most species in these two genera are related to intestinal diseases [2]. Hence, this biosensor platform is an efficient tool for culture-independent, rapid, ultrasensitive, and discriminative detection of *Campylobacter* and *Aliarcobacter* genera. The specificity for different species in *Campylobacter* or *Aliarcobacter* in biosensing can hopefully be achieved via species-specific aptamers to develop novel species-discriminative biosensors.

## 4. Conclusion

In conclusion, the microfluidics-assisted biosensing platform shown here offers culture-independent, rapid, ultra-sensitive, and discriminative detection of *Campylobacter* and *Aliarcobacter* in human stool and chicken samples. The advances of this platform are attributed to the properties of its modules including the general capture of different species of the FsC[Fe^3+^]-modified MNPs, genera-selectivity of the aptamer-modified CDs, and neglectable interference along with the extraordinary luminescent properties of the red- and green-emitting CDs that render the ultra-sensitivity. Exclusively, this work first demonstrates that the fungal siderophore FsC can serve as a ligand for *Campylobacter* and *Aliarcobacter* to be used to synthesize a new magnetic material for selective capture of bacteria. Moreover, this work suggests that a visual RGB indication, which is transformed from the fluorescence signals of the twin CDs-enabled dual-fluorescent detection channels, is able to aid the differentiation between the two genera. This approach can be extended to the discrimination of other close species in the same genus or even different serotypes from the same species This would be achieved by the discovery of new ligands for fabricating selective capture modules, selection of species- or serotype-specific aptamers, and the development of novel triple- or multiple-channel fluorescent/colorimetric transducing methodologies. With them integrated as novel biosensors, the RGB discrimination for multiple bacteria by using a portable device, such as a smartphone, is hopeful to be achieved.

## Acknowledgements

This work was supported by the Bingtuan Science and Technology Program [Grant No. 2021BC009]. We also thank Professor Maojun Zhang from Chinese Center for Disease Control and Prevention for her kind directions about *Camplylobacter* and *Aliarcobacter*. We thank LetPub (www.letpub.com) for linguistic assistance and pre-submission expert review.

## Appendix A Supplementary material

Supplementary data associated with this article can be found in the online version.

