## Supplemental Figures, Tables and Methods for "A biosensor encompassing fusarinine C-magnetic nanoparticles and aptamer-red/green carbon dots for dual-channel fluorescent and RGB discrimination of *Campylobacter* and *Aliarcobacter*"

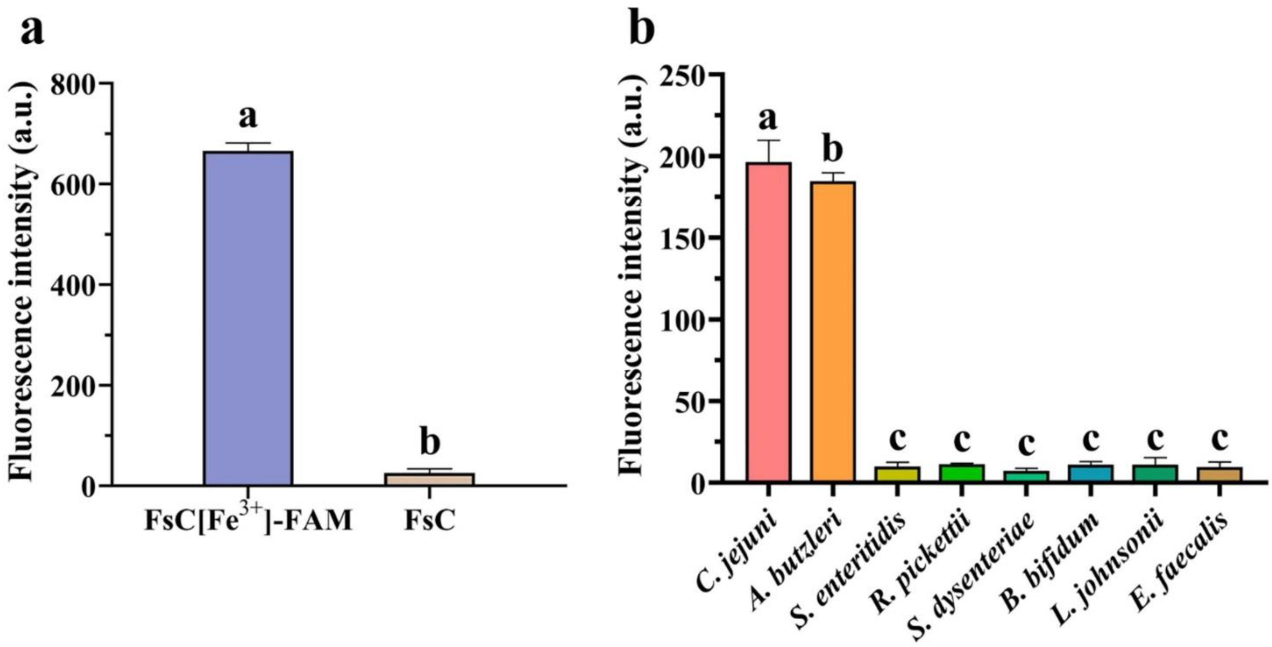


**Fig. S1.** (a) Fluorescence intensities of 5-FAM modified FsC[$Fe^{3+}$] and FsC[$Fe^{3+}$] to verify the successful conjugation between 5-FAM and FsC[$Fe^{3+}$]. (b) Active targeting of FsC[$Fe^{3+}$] to other intestinal bacteria compared with those for *C. jejuni* and *A. butzleri* jusitified by using the FsC[$Fe^{3+}$]-FAM. λEx: 493 nm, λEm: 522 nm. Data are presented as mean ± SD, n= 3. abc, data with different symbols have significant difference.

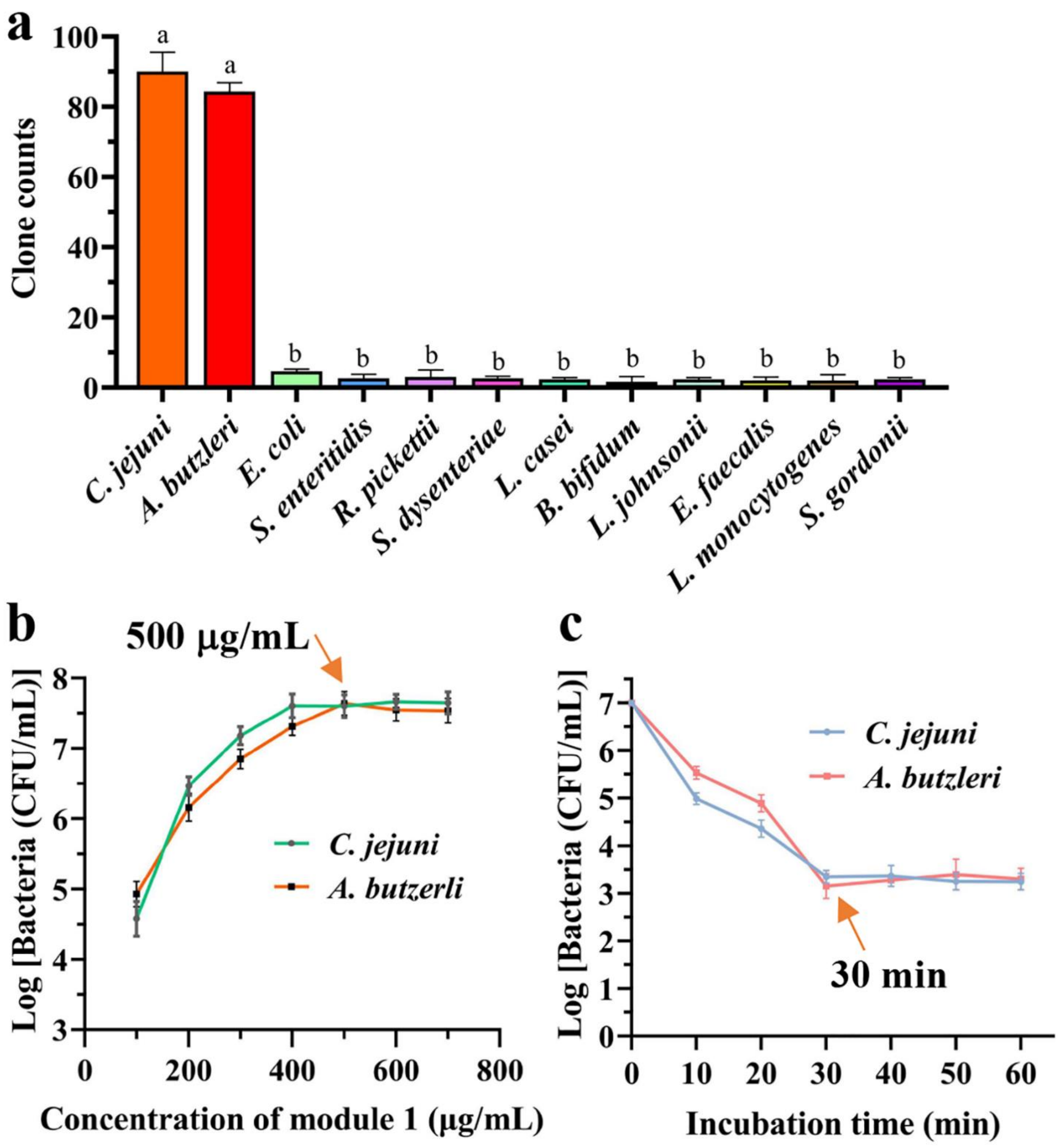


**Fig. S2.** (a) Analysis of 16S rDNA tag to count the number of different bacteria captured by module 1 (Ca/FsC[$Fe^{3+}$]-MNPs) from the artificial bacterial community. (b) Optimization of module 1 concentration for capturing *C. jejuni* or *A. butzleri*. The optimal concentration was verified to be 500 μg/mL marked with a red cycle. (c) Optimization of reacting time between module 1 and *C. jejuni* or *A. butzleri*. The optimal time was verified to be 30 min marked with a green box. Data are presented as mean ± SD, n= 3. ab, data with different symbols have significant differences.

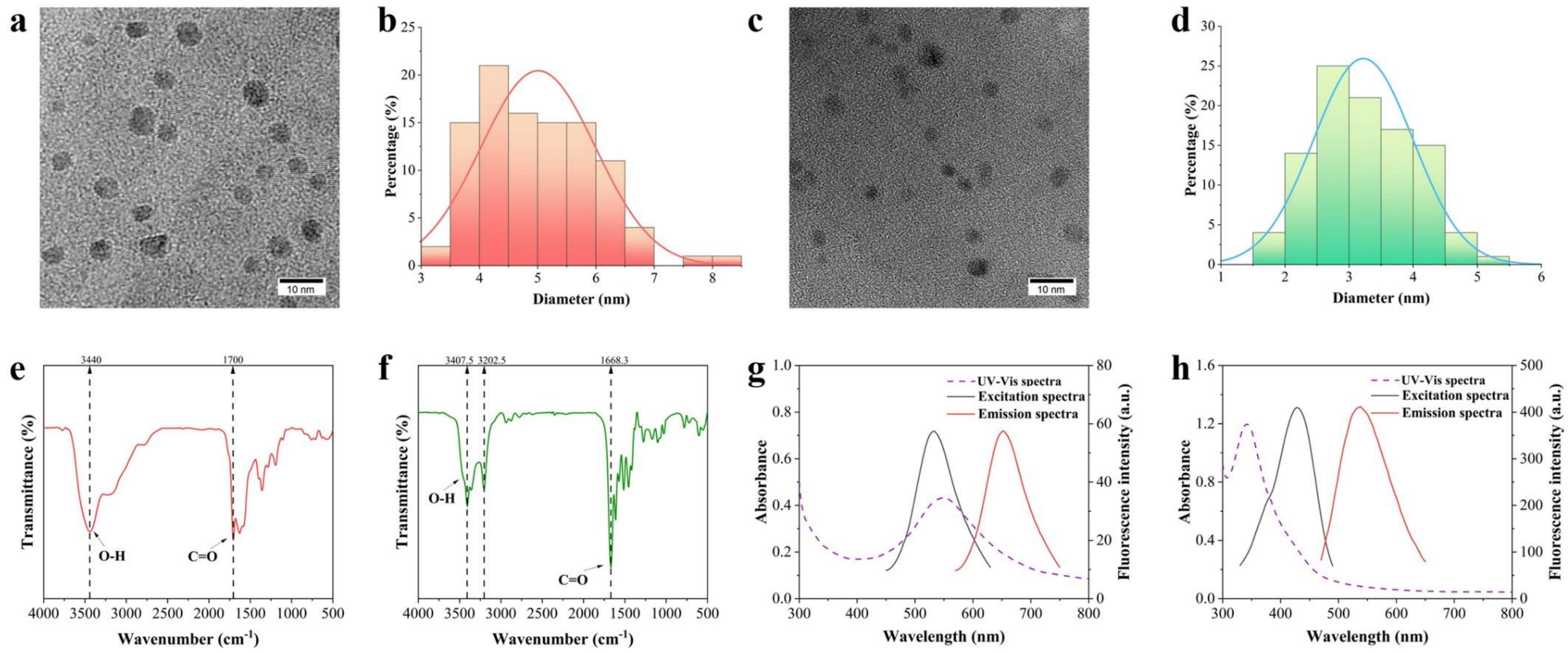


**Fig. S3.** Characterizations of carbon dots (CDs). (a, b, c, d) TEM iamges and average particle sizes of red- and green-emitting CDs (RCDs and GCDs). Scale bar: 10.0 nm for RCDs, 5.0 nm for GCDs. (c, d) FT-IR spectra of RCDs and GCDs. (e, f) UV-Vis spectra and fluorescence spectra of RCDs and GCDs.

In **Fig. S3a and b**, it could be seen that RCDs were evenly distributed with an average particle size of approximately 5.0 nm. Visually, the GCDs were smaller in size than that of RCDs (**Fig. S3c**) observed under the same measuring scale of 10.0 nm, with an average particle size of approximately 3 nm (**Fig. S3d**). The FT-IR spectra of RCDs (**Fig. S3c**)indicated characteristic peaks at 1700 $cm^{-1}$ and 3440 $cm^{-1}$, which were ascribed to C=O and O-H bond vibrations in the spectrum, respectively [1]. The FT-IR spectra of GCDs (**Fig. S3d**) showed characteristic peaks at 1668.3 $cm^{-1}$ and 3407.5 $cm^{-1}$, which were ascribed to C=O and O-H bond vibrations in the spectrum, respectively [2]. These results were in consistence with the previous work [1, 2], indicating the harboring of carboxyl groups on the surface of the above two CDs. Moreover, the purple dotted line representing the UV-Vis spectrum in **Fig. S3e** indicated the maximal absorption peak was at 556.0 nm for RCDs. The black and red lines represented the excitation and emission spectra, respectively. At the excitation wavelength of 545.0 nm, the maximum emission wavelength was at 640.0 nm. Likewise, **Fig. S3f** showed the maximal absorption peak of GCDs was at 340.0 nm in the UV-Vis spectrum; the maximal emission wavelength was 420.0 nm with the maximal excitation wavelength at 530.0 nm. These data were also identical to the referenced work [1, 2], justifying that RCDs and GCDs were correctly synthesized.

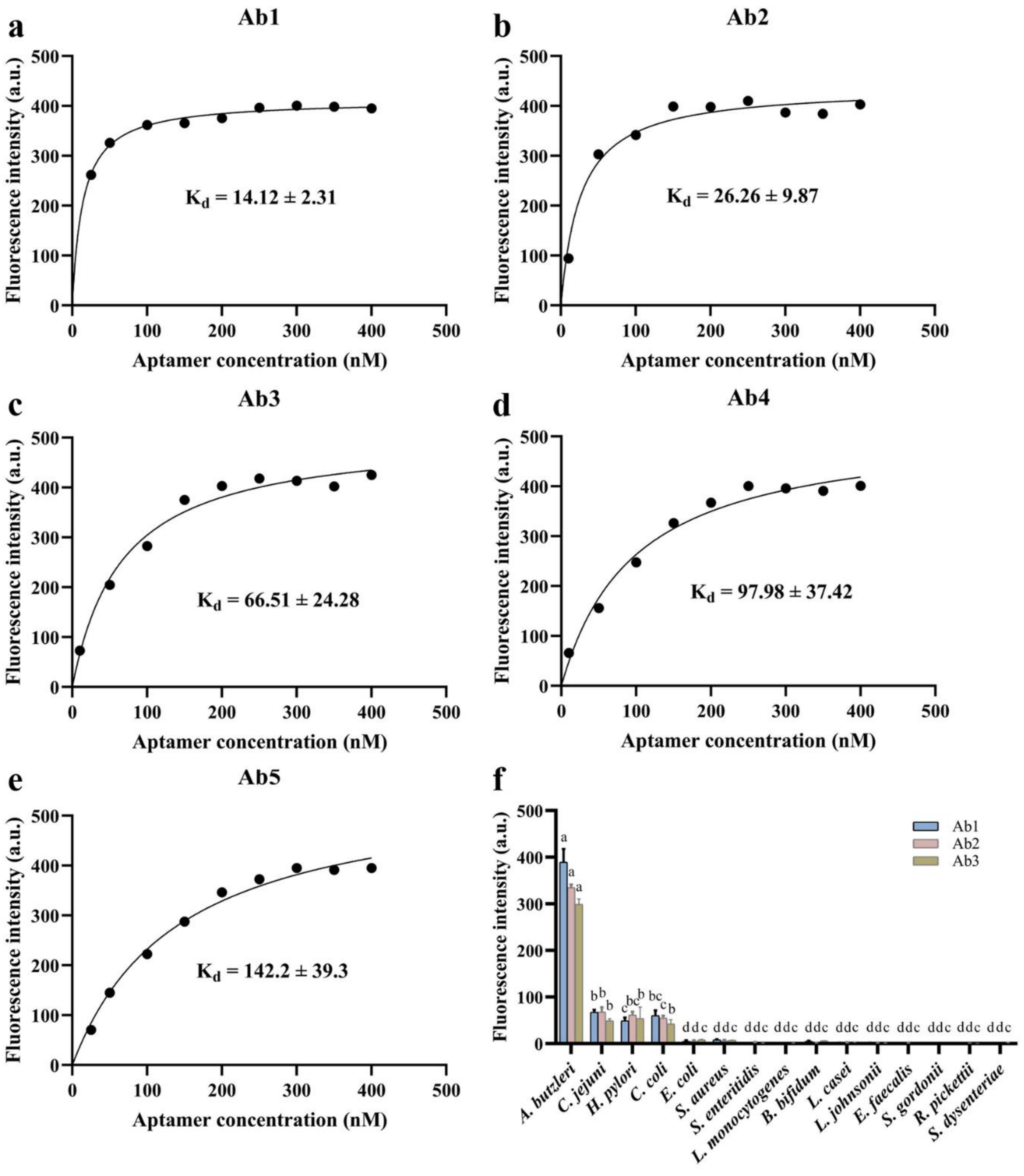


**Fig. S4.** (a–e) Equilibrium dissociation constant (Kd) representing the binding affinity of each aptamer candidate to *A. butzleri* cells determined by using the synthesized FAM–aptamer. (f) Specificity of candidate aptamers Ab1, Ab2, and Ab3 to *A. butzleri* identified using the corresponding FAM–aptamers. λEx: 493 nm, λEm: 522 nm. Data, where necessary, are presented as mean ± SD, n = 3. abcd, data with different symbols have significant difference.

Ab1, Ab2, Ab3, Ab4, Ab5 were the five sequences with the highest abundance in the ssDNA pool from the 12th round of SELEX identified by high-throughput sequencing; the strand numbers for them were 2295158 (Ab1), 295827 (Ab2), 294330 (Ab3), 285805 (Ab4), 263782 (Ab5), respectively. As revealed by their Kd values, the aptamers of Ab1, Ab2, and Ab3 had high binding affinities, and they were selected to be tested for theri specificities for *A. butzleri*. With the synthesize FAM–5'–aptamer probes for these three aptamers, Ab1 was eventually recognized as the best aptamer with the highest specificity to *A. butzleri*. The nucleic acid sequence of the candidate aptamers (Ab1, Ab2, Ab3, Ab4 and Ab5) were as follows:

Ab1:

TGGTGCGTGCTATTCAGAGTGAGGGACGCATGATTGGGTGACTCCGGGAGATCATGCAAGGAACCTGT

AGCCACGAATAC

Ab2:

TGGTGCGTGCTATTCAGAGTGGGGTGGACCGGGGGGATCGAATAACGCATCGCGCCTAGTGAACCTGT

AGCCACGAATAC

Ab3:

TGGTGCGTGCTATTCAGAGTCCTCTAGGCGGTAAGGATCGGACCTTGGTCTGATGGTGAGGAACCTGTA

GCCACGAATAC

Ab4:

TGGTGCGTGCTATTCAGAGTGACGGATACGGATTCAGGGCGGAGTGCCAAGATACGGATGGAACCTGT

AGCCACGAATAC

Ab5:

TGGTGCGTGCTATTCAGAGTGTTGAGGATTAGGATCGCTTTAAACGATTAGGATCCGAATGAACCTGTA

GCCACGAATAC

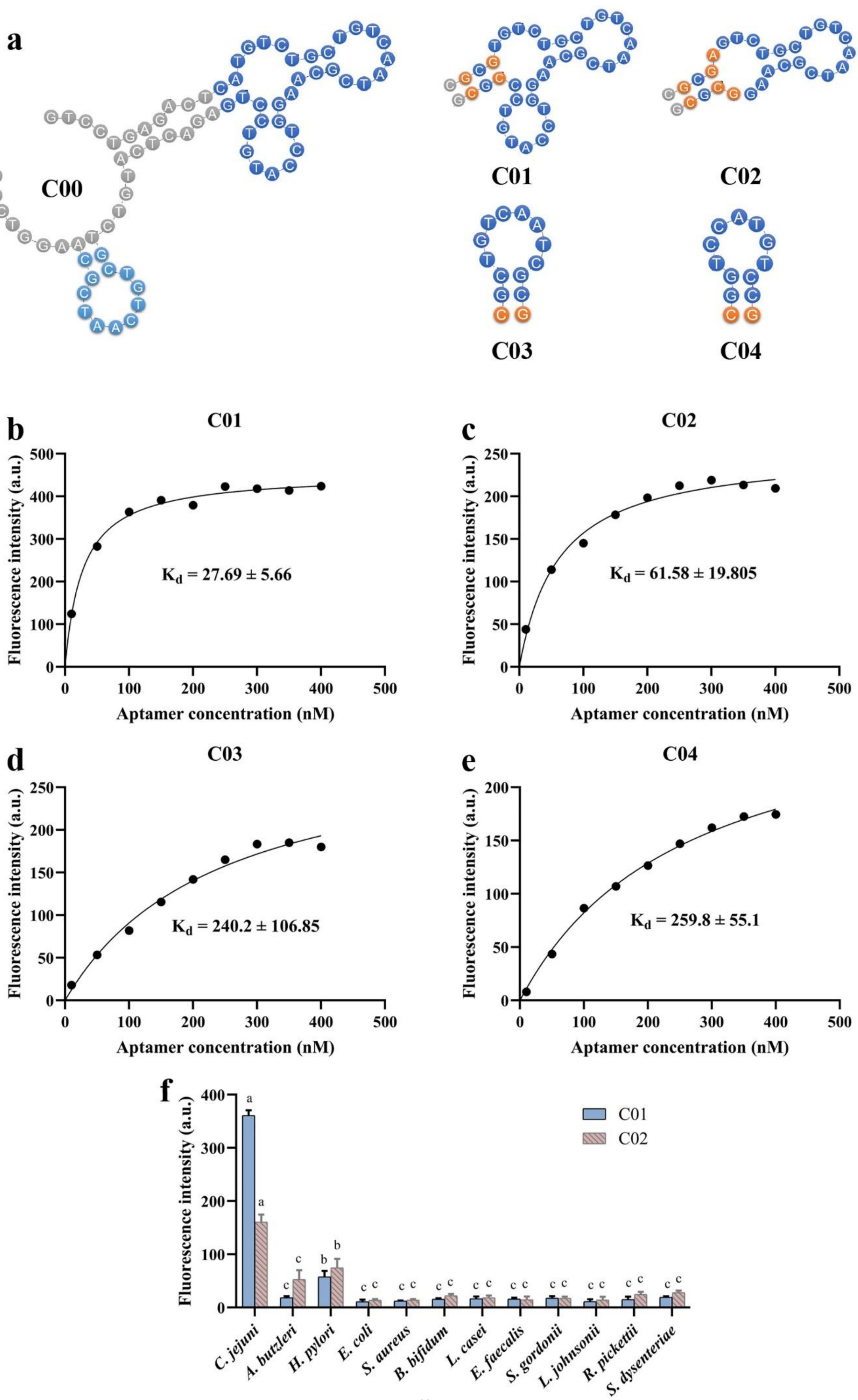
a
C00
C01
C02
C03
C04
b
C01
Fluorescence intensity (a.u.)
Aptamer concentration (nM)
K$_d$ = 27.69 ± 5.66
c
C02
Fluorescence intensity (a.u.)
Aptamer concentration (nM)
K$_d$ = 61.58 ± 19.805
d
C03
Fluorescence intensity (a.u.)
Aptamer concentration (nM)
K$_d$ = 240.2 ± 106.85
e
C04
Fluorescence intensity (a.u.)
Aptamer concentration (nM)
K$_d$ = 259.8 ± 55.1
f
Fluorescence intensity (a.u.)
C01
C02
C. jejuni
A. butzleri
H. pylori
E. coli
S. aureus
B. bifidum
L. casei
E. faecalis
S. gordonii
L. johnsonii
R. pickettii
S. dysenteriae

**Fig. S5.** (a) Graphical illustration for the truncation of the *C. jejuni*-specific aptamer. (b-e) Equilibrium dissociation constants (Kd) reflecting the binding affinity of each truncated aptamer candidate to *C. jejuni* cells determined using the corresponding FAM–aptamer probes. (f) Specificity of truncated aptamers C01 and C02 to *C. jejuni* identified using the corresponding FAM–aptamer probes. λEx: 493 nm, λEm: 522 nm. Data, where necessary, are presented as mean ± SD, n = 3. abc, data with different symbols have significant difference.

As shown in **Fig. S5a**, the secondary structures of the original *C. jejuni*-specific aptamer (C00) and the truncated ones (C01, C02, C03 and C04) were predicted by DNAMAN. The 4 truncated aptamers were obtained by reasonably intercepting and appropriately replacing some bases. Among them, C01 was the main region of the secondary structure of C00. Furthermore, based on C01, the minor ring species were intercepted to obtain C02, C03 and C04 aptamer. **Fig. S5b-e** showed the binding curves of the 4 truncated aptamers with Kd values of 27.69 ± 5.66 nM, 61.58 ± 19.805 nM, 240.2 ± 106.85 nM and 259.8 ± 55.1 nM, respectively. Among them, the Kd value of C01 was the lowest, indicating its highest affinity. Moreover, compared with C00 (Kd=48.12 ± 6.15 nM), its affinity was nearly doubled. In **Fig. S5f**, it was demonstrated that C01 had significantly high specificity to *C. jejuni* than that of C02. Therefore, C01 was eventually recognized as the most appropriate truncated aptamer specific for *C. jejuni*, and it was used to synthesize ARCDs. The nucleic acid sequences of the original [3] and truncated aptamers were as follows:

C00:

GTCCTGAGACTCATGTCTGCTGTCAATCGCAAGGTCCATGTCCTGAGACTCATGTCTGCTGTCAATCGC

AAGGTCCA

C01: CGCGTGTCTGCTGTCAATCGCAAGGTCCATGTCCCGCG

C02: CGCGAGTCTGCTGTCAATCGCAAGGCGCG

C03: CGCTGTCAATCGCG

C04: GCGTCCATGTCGC

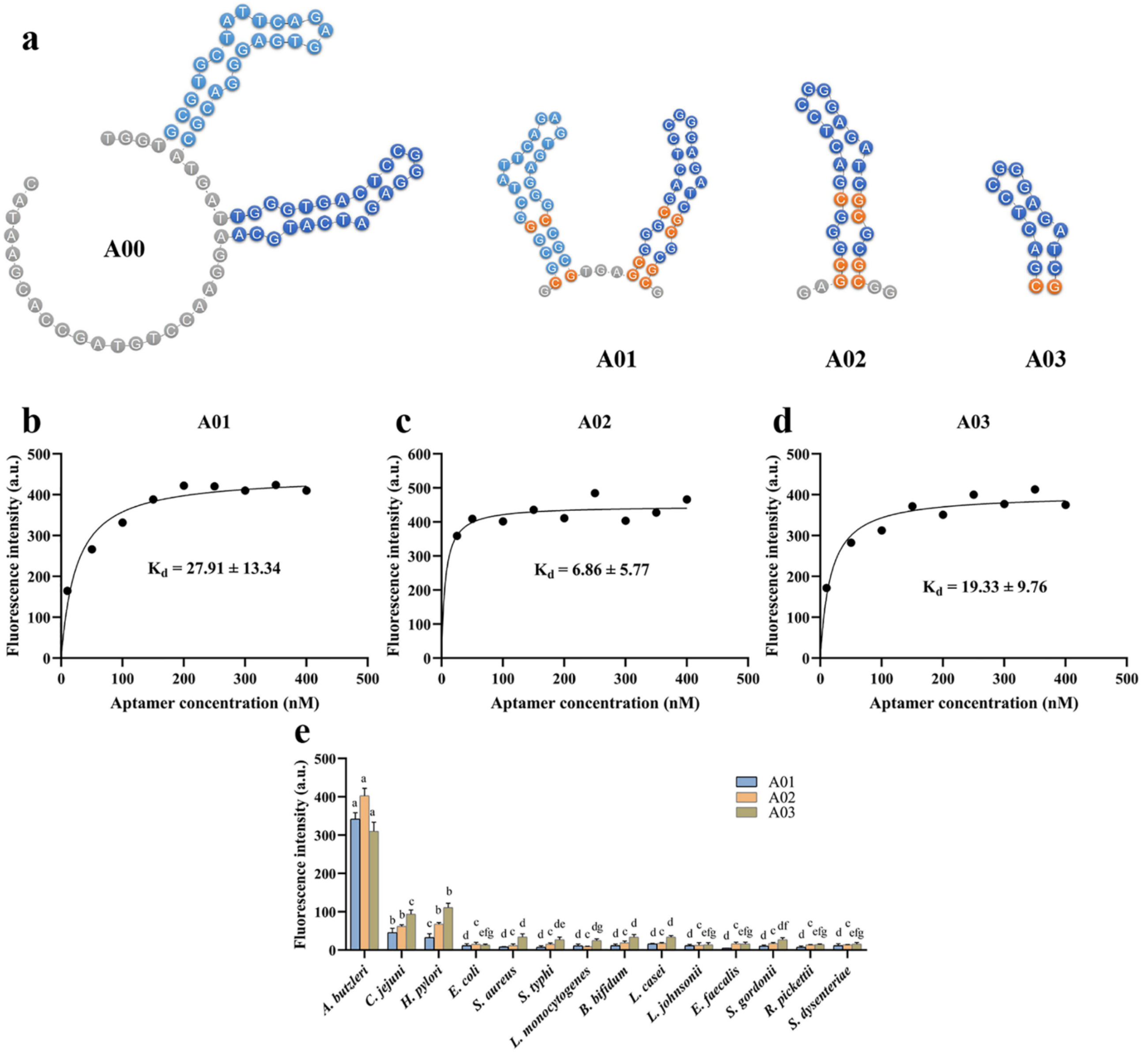


**Fig. S6.** (a) Graphical illustration for the truncation of the *A. butzleri*-specific aptamer. (b-d) Equilibrium dissociation constants (Kd) reflecting the binding affinity of each truncated aptamer candidate to *A. butzleri* cells determined using the corresponding FAM–aptamer probes. (e) Specificity of truncated aptamers A01, A02 and A03 to *A. butzleri* identified using the corresponding FAM–aptamer probes. λEx: 493 nm, λEm: 522 nm. Data, where necessary, are presented as mean ± SD, n = 3. abcdefg, data with different symbols have significant difference.

Similarly, as shown in **Fig. S6a**, the secondary structures of the original *A. butzleri*-specific aptamer (A00) and the truncated ones (A01, A02 and A03) were predicted by DNAMAN. The principle and method of truncation were consistent with the above. Briefly, A01, the main region of the secondary structure of A00, contained two stem-loop structures. The stem-loop structure of A02 without primer sequence intercepted a valid sequence from A01. Further, aptamer A03 was intercepted from A02 to obtain a shorter aptamer. The highlighted base in the truncated aptamer structure was the replaced base to maintain the stability of the secondary structure. **Fig. S6b-d** showed the binding curves of 3 truncated aptamers with Kd values of 27.91 ± 13.34 nM, 6.86 ± 5.77 nM and 19.33 ± 9.76 nM, respectively. Among them, the Kd value of A02 was the lowest, indicating its highest affinity. Furthermore, compared with A00 (Kd = 14.12 ± 2.31 nM), its affinity was greatly improved, while the affinity of A01 and A03 were inconspicuously changed. **Fig. S6e** showed that the specificity of A02 was not significantly different from A01 and slightly better than A03. Thus, A02 was eventually recognized as the most appropriate truncated aptamer specific for *A. butzleri*, and it was used to synthesize AGCDs. The nucleic acid sequence of the original and truncated aptamers were as follows:

A00:

TGGTGCGTGCTATTCAGAGTGAGGGACGCATGATTGGGTGACTCCGGGAGATCATGCAAGGAACCTGT

AGCCACGAATAC

A01: GCGCGGGCTATTCAGAGTGAGGGCCGCGTGAGCGGGCGACTCCGGGAGATCGCGCGCG

A02: GAGCGGGCGACTCCGGGAGATCGCGCGCGG

A03: CGACTCCGGGAGATCG

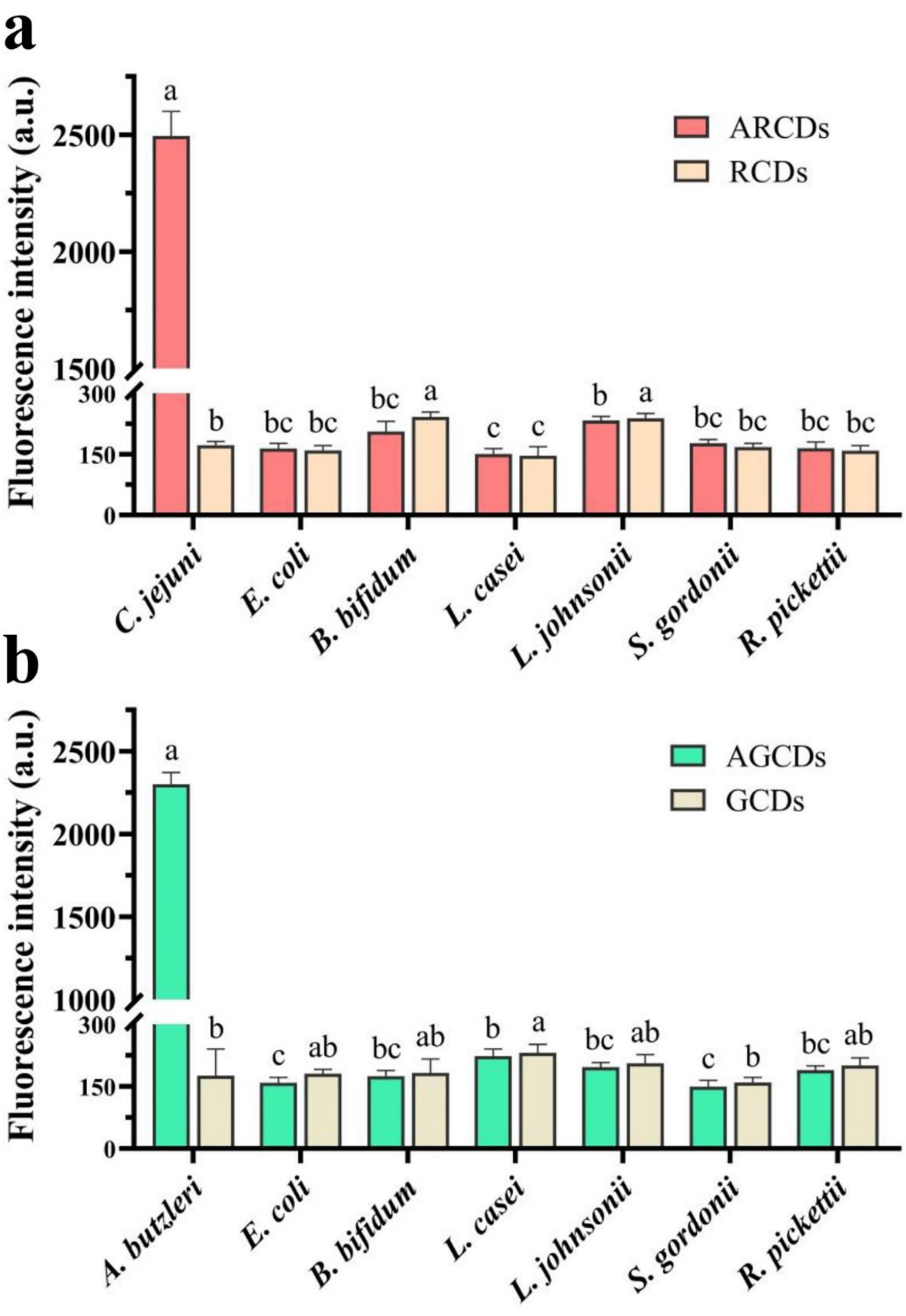


**Fig. S7.** The validated specificity of ARCDs for *C. jejuni* (a) and AGCDs for *A. butzleri* (b) against other intestinal bacteria, manifested as significantly higher emitted fluorescence intensities than those of the control bacteria after the CDs were incubated with each bacterium accordingly. Data are presented as mean ± SD, n = 3. abc, data with different symbols have significant difference.

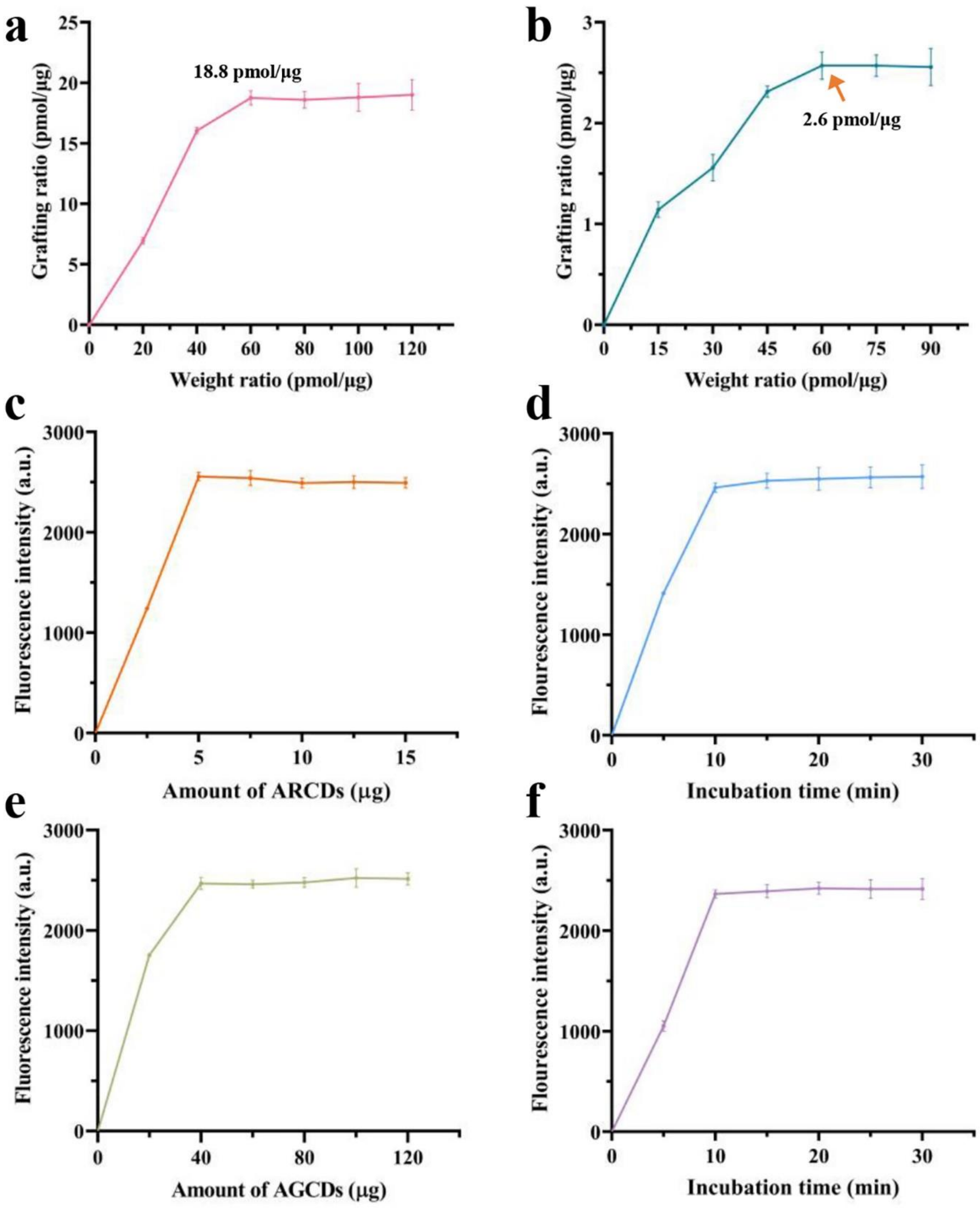


**Fig. S8.** Optimizations of ARCDs and AGCDs in module 2. (a, b) Optimal mole/weight ratio between aptamers and CDs for obtaining ARCDs and AGCDs with the highest aptamer grafting ratios. (c, d) Optimal weight and incubation time for ARCDs to indicate $1.0 \times 10^7$ CFU/mL *C. jejuni*. (e, f) Optimal weight and incubation time for AGCDs to indicate $1.0 \times 10^7$ CFU/mL *A. butzleri*. Data are presented as mean ± SD, n = 3.

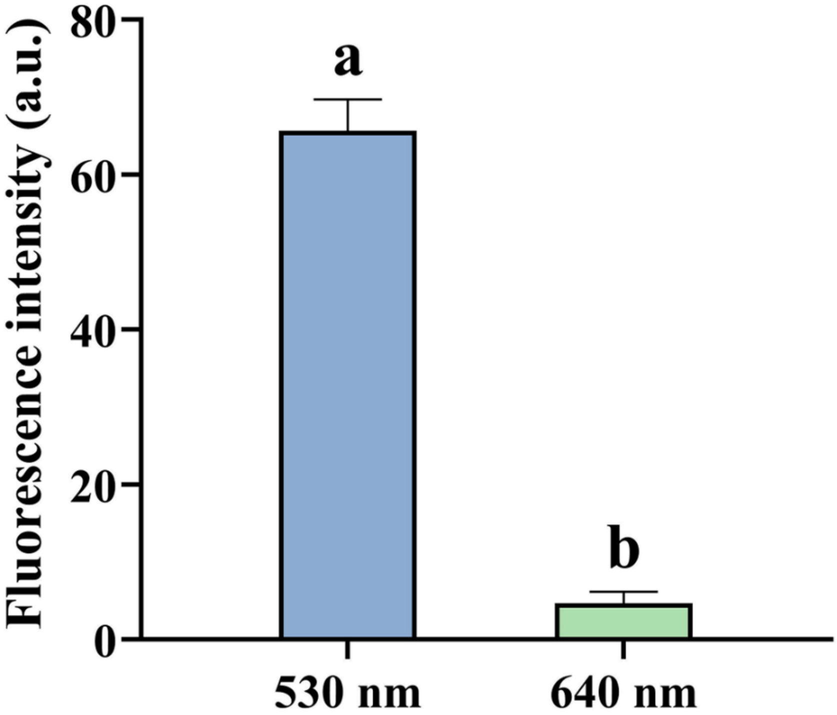


**Fig. S9.** Intensities of emitted fluorescence from module 1 under the excitation wavelength of AGCDs (530.0 nm) and ARCDs (640.0 nm). ab, data with different symbols have significant difference.

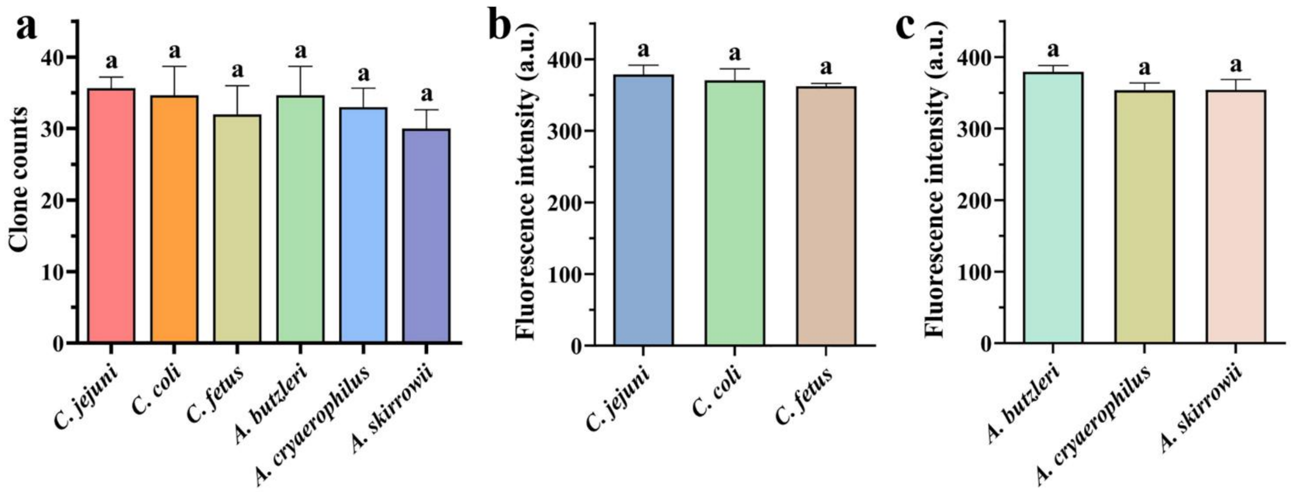


**Fig. S10.** Selectivities of the biosensor modules for different species of *Campylobacter* and *Aliarcobacter*. (a) Capturing efficiencies of module 1 for different species of *Campylobacter* and *Aliarcobacter* determined by the clone counts of each species captured by module 1. (b, c) aptamer C01 and aptamer A02 for different species of *Campylobacter* and *Aliarcobacter*. a, data with the same symbol are not significantly different.

**Table S1.** Microfluidic collected *C. jejuni* and *A. butzleri* in simulated chicken meat samples.

| *A. butzleri* (CFU/mL) / *C. jejuni* (CFU/mL) | 0 | $1\times10^{1}$ | $1\times10^{2}$ | $1\times10^{3}$ | $1\times10^{4}$ |
|---|---|---|---|---|---|
| 0 | $0^{c}$, $0^{c}$ | $0^{c}$, $6.3 \pm 0.6^{c}$ | $0^{c}$, $66.7 \pm 4.5^{c}$ | $0^{c}$, $655.2 \pm 15.6^{b}$ | $0^{c}$, $(6.4 \pm 0.2)\times10^{3\ a}$ |
| $1\times10^{1}$ | $6.4 \pm 1.2^{c}$, $0^{c}$ | $5.4 \pm 0.5^{c}$, $5.7 \pm 0.6^{c}$ | $5.4 \pm 1.5^{c}$, $55.8 \pm 2.9^{c}$ | $4.3 \pm 0.6^{c}$, $594.2 \pm 57.2^{b}$ | $5.2 \pm 0.5^{c}$, $(6.5 \pm 0.2)\times10^{3\ a}$ |
| $1\times10^{2}$ | $57.2 \pm 6.1^{c}$, $0^{c}$ | $55.1 \pm 7.1^{c}$, $4.3 \pm 0.6^{c}$ | $53.9 \pm 7.0^{c}$, $51.7 \pm 6.6^{c}$ | $54.3 \pm 3.0^{c}$, $608.6 \pm 23.6^{b}$ | $51.6 \pm 4.8^{c}$, $(6.2 \pm 0.2)\times10^{3\ a}$ |
| $1\times10^{3}$ | $652.5 \pm 26.6^{b}$, $0^{c}$ | $629.1 \pm 18.6^{b}$, $4.3 \pm 0.6^{c}$ | $570.0 \pm 20.4^{b}$, $48.3 \pm 3.3^{c}$ | $539.3 \pm 28.9^{b}$, $570.9 \pm 14.5^{b}$ | $524.8 \pm 52.6^{b}$, $(5.8 \pm 0.2)\times10^{3\ a}$ |
| $1\times10^{4}$ | $(6.7 \pm 0.4)\times10^{3\ a}$, $0^{c}$ | $(6.3 \pm 0.3)\times10^{3\ a}$, $4.4 \pm 0.2^{c}$ | $(5.9 \pm 0.2)\times10^{3\ a}$, $50.6 \pm 1.8^{c}$ | $(5.5 \pm 0.3)\times10^{3\ a}$, $562.4 \pm 30.0^{b}$ | $(5.3 \pm 0.3)\times10^{3\ a}$, $(5.5 \pm 0.3)\times10^{3\ a}$ |

Data are presented as mean ± SD, n = 3. abc, data with different symbols are significantly different.

**Table S2.** Microfluidic collected *C. jejuni* and *A. butzleri* in simulated fecal samples.

| *C. jejuni* (CFU/mL) \ *A. butzleri* (CFU/mL) | 0 | $1\times10^{1}$ | $1\times10^{2}$ | $1\times10^{3}$ | $1\times10^{4}$ | $1\times10^{5}$ | $1\times10^{6}$ | $1\times10^{7}$ |
|---|---|---|---|---|---|---|---|---|
| 0 | $0^{c}, 0^{d}$ | $0^{d}, 6.7 \pm 0.6^{d}$ | $0^{c}, 67.3 \pm 3.2^{d}$ | $0^{c}, 669.2 \pm 16.3^{d}$ | $0^{c}, (6.5 \pm 0.1)\times10^{3\ d}$ | $0^{c}, (6.6 \pm 0.2)\times10^{4\ c}$ | $0^{c}, (6.4 \pm 0.2)\times10^{5\ b}$ | $0^{c}, (6.2 \pm 0.1)\times10^{6\ a}$ |
| $1\times10^{1}$ | $5.7 \pm 1.5^{c}, 0^{c}$ | $5.4 \pm 0.5^{d}, 5.7 \pm 0.6^{c}$ | $5.2 \pm 0.2^{c}, 55.2 \pm 2.1^{c}$ | $4.3 \pm 0.6^{c}, 638.5 \pm 32.6^{c}$ | $3.8 \pm 0.1^{c}, (6.2 \pm 0.1)\times10^{3\ c}$ | $5.3 \pm 1.7^{c}, (6.4 \pm 0.1)\times10^{4\ c}$ | $4.8 \pm 1.1^{c}, (6.2 \pm 0.2)\times10^{5\ b}$ | $3.9 \pm 0.4^{c}, (5.8 \pm 0.3)\times10^{6\ a}$ |
| $1\times10^{2}$ | $65.4 \pm 6.0^{c}, 0^{c}$ | $57.2 \pm 2.1^{d}, 4.7 \pm 0.6^{c}$ | $49.6 \pm 3.7^{c}, 55.6 \pm 2.7^{c}$ | $52.7 \pm 1.9^{c}, 632.0 \pm 38.3^{c}$ | $54.0 \pm 2.8^{c}, (6.3 \pm 0.3)\times10^{3\ c}$ | $52.0 \pm 2.1^{c}, (6.3 \pm 0.2)\times10^{4\ c}$ | $50.9 \pm 4.0^{c}, (6.0 \pm 0.1)\times10^{5\ b}$ | $51.2 \pm 1.3^{c}, (5.8 \pm 0.2)\times10^{6\ a}$ |
| $1\times10^{3}$ | $654.4 \pm 11.3^{c}, 0^{c}$ | $624.9 \pm 15.6^{d}, 4.7 \pm 1.2^{c}$ | $593.3 \pm 32.4^{c}, 49.9 \pm 5.1^{c}$ | $539.3 \pm 28.9^{c}, 570.9 \pm 14.5^{c}$ | $551.9 \pm 31.3^{c}, (6.1 \pm 0.1)\times10^{3\ c}$ | $525.3 \pm 28.5^{c}, (6.2 \pm 0.2)\times10^{4\ c}$ | $518.0 \pm 15.6^{c}, (5.7 \pm 0.1)\times10^{5\ b}$ | $499.7 \pm 18.9^{c}, (5.8 \pm 0.3)\times10^{6\ a}$ |
| $1\times10^{4}$ | $(6.3 \pm 0.1)\times10^{3\ c}, 0^{c}$ | $(6.4 \pm 0.1)\times10^{3\ d}, 5.5 \pm 0.2^{c}$ | $(5.9 \pm 0.2)\times10^{3\ c}, 47.7 \pm 2.3^{\ c}$ | $(5.4 \pm 0.1)\times10^{3\ c}, 614.8 \pm 14.6^{c}$ | $(5.5 \pm 0.2)\times10^{3\ c}, (6.2 \pm 0.1)\times10^{3\ c}$ | $(5.4 \pm 0.1)\times10^{3\ c}, (6.0 \pm 0.1)\times10^{4\ c}$ | $(5.7 \pm 0.2)\times10^{3\ c}, (5.8 \pm 0.1)\times10^{5\ b}$ | $(5.4 \pm 0.3)\times10^{3\ c}, (5.6 \pm 0.2)\times10^{6\ a}$ |
| $1\times10^{5}$ | $(6.3 \pm 0.1)\times10^{4\ c}, 0^{c}$ | $(6.2 \pm 0.2)\times10^{4\ c}, 4.9 \pm 0.8^{c}$ | $(6.0 \pm 0.1)\times10^{4\ c}, 50.5 \pm 3.2^{c}$ | $(5.8 \pm 0.1)\times10^{4\ c}, 594.0 \pm 7.2^{c}$ | $(5.8 \pm 0.1)\times10^{4\ c}, (5.7 \pm 0.1)\times10^{3\ c}$ | $(5.6 \pm 0.1)\times10^{4\ c}, (5.7 \pm 0.3)\times10^{4\ c}$ | $(5.4 \pm 0.2)\times10^{4\ c}, (5.4 \pm 0.1)\times10^{5\ b}$ | $(4.9 \pm 0.2)\times10^{4\ c}, (5.4 \pm 0.2)\times10^{6\ a}$ |
| $1\times10^{6}$ | $(6.5 \pm 0.3)\times10^{5\ b}, 0^{d}$ | $(6.2 \pm 0.1)\times10^{5\ b}, 5.3 \pm 0.6^{d}$ | $(6.0 \pm 0.1)\times10^{5\ b}, 51.2 \pm 1.5^{d}$ | $(5.7 \pm 0.2)\times10^{5\ b}, 569.5 \pm 15.9^{d}$ | $(5.6 \pm 0.2)\times10^{5\ b}, (5.5 \pm 0.2)\times10^{3\ cd}$ | $(5.4 \pm 0.2)\times10^{5\ b}, (5.4 \pm 0.1)\times10^{4\ c}$ | $(5.2 \pm 0.2)\times10^{5\ b}, (5.3 \pm 0.1)\times10^{5\ b}$ | $(4.9 \pm 0.2)\times10^{5\ b}, (5.3 \pm 0.1)\times10^{6\ a}$ |
| $1\times10^{7}$ | $(6.4 \pm 0.2)\times10^{6\ a}, 0^{d}$ | $(6.3 \pm 0.1)\times10^{6\ a}, 4.5 \pm 0.7^{d}$ | $(5.6 \pm 0.2)\times10^{6\ a}, 49.5 \pm 4.7^{d}$ | $(5.0 \pm 0.1)\times10^{6\ a}, 530.3 \pm 19.7^{\ cd}$ | $(4.8 \pm 0.1)\times10^{6\ a}, (5.2 \pm 0.3)\times10^{3\ cd}$ | $(4.9 \pm 0.1)\times10^{6\ a}, (5.1 \pm 0.1)\times10^{4\ c}$ | $(4.9 \pm 0.2)\times10^{6\ a}, (5.1 \pm 0.2)\times10^{5\ b}$ | $(5.0 \pm 0.2)\times10^{6\ a}, (4.9 \pm 0.1)\times10^{6\ a}$ |

Data are presented as mean ± SD, n = 3. abcd, data with different symbols are significantly different.

**Table S3.** Detection of *C. jejuni* and *A. butzleri* in simulated chicken meat samples by dual-channel biosensor.

| *A. butzleri* (CFU/mL) / *C. jejuni* (CFU/mL) | 0 | $1\times10^1$ | $1\times10^2$ | $1\times10^3$ | $1\times10^4$ |
|---|---|---|---|---|---|
| 0 | $0^c$, $0^c$ | $0^c$, $5.7 \pm 0.5^c$ | $0^c$, $61.5 \pm 5.0^c$ | $0^c$, $644.6 \pm 12.3^b$ | $0^c$, $(6.3 \pm 0.4)\times10^{3\ a}$ |
| $1\times10^1$ | $5.4 \pm 1.6^c$, $0^c$ | $3.8 \pm 1.5^c$, $4.8 \pm 0.9^c$ | $5.0 \pm 0.9^c$, $53.0 \pm 2.3^c$ | $4.0 \pm 1.1^c$, $556.2 \pm 11.2^b$ | $4.2 \pm 1.1^c$, $(6.2 \pm 0.3)\times10^{3\ a}$ |
| $1\times10^2$ | $52.3 \pm 5.3^c$, $0^c$ | $52.4 \pm 6.9^c$, $4.3 \pm 0.9^c$ | $50.3 \pm 6.1^c$, $48.0 \pm 6.3^c$ | $48.2 \pm 4.0^c$, $596.1 \pm 23.3^b$ | $46.3 \pm 5.7^c$, $(6.0 \pm 0.6)\times10^{3\ a}$ |
| $1\times10^3$ | $639.5 \pm 19.9^b$, $0^c$ | $609 \pm 7.3^b$, $3.6 \pm 0.4^c$ | $544.9 \pm 11.6^b$, $45.4 \pm 2.1^c$ | $507.4 \pm 25.2^b$, $556.2 \pm 11.2^b$ | $511.2 \pm 29.6^b$, $(5.9 \pm 0.6)\times10^{3\ a}$ |
| $1\times10^4$ | $(6.4 \pm 0.4)\times10^{3\ a}$, $0^c$ | $(6.0 \pm 0.3)\times10^{3\ a}$, $4.1 \pm 2.1^c$ | $(5.6 \pm 0.5)\times10^{3\ a}$, $48.3 \pm 10.2^c$ | $(5.2 \pm 0.2)\times10^{3\ a}$, $531.8 \pm 50.5^b$ | $(5.0 \pm 0.3)\times10^{3\ a}$, $(5.6 \pm 0.2)\times10^{3\ a}$ |

Data are presented as mean ± SD, n = 3. abc, data with different symbols are significantly different.

**Table S4.** Detection of *C. jejuni* and *A. butzleri* in simulated fecal samples by dual-channel biosensor

| *C. jejuni* (CFU/mL) \ *A. butzleri* (CFU/mL) | 0 | $1\times10^{1}$ | $1\times10^{2}$ | $1\times10^{3}$ | $1\times10^{4}$ | $1\times10^{5}$ | $1\times10^{6}$ | $1\times10^{7}$ |
|---|---|---|---|---|---|---|---|---|
| 0 | $0^{c}, 0^{d}$ | $0^{d}, 6.0 \pm 1.1^{d}$ | $0^{c}, 62.9 \pm 2.6^{d}$ | $0^{d}, 651.8 \pm 21.6^{d}$ | $0^{c}, (6.5 \pm 0.3)\times10^{3\ cd}$ | $0^{d}, (6.4 \pm 0.3)\times10^{4\ c}$ | $0^{c}, (6.3 \pm 0.2)\times10^{5\ b}$ | $0^{c}, (6.1 \pm 0.1)\times10^{6\ a}$ |
| $1\times10^{1}$ | $4.9 \pm 1.7^{c}, 0^{d}$ | $4.6 \pm 1.1^{d}, 5.7 \pm 0.5^{d}$ | $4.7 \pm 0.6^{c}, 49.1 \pm 2.2^{d}$ | $3.6 \pm 1.3^{d}, 623.6 \pm 24.3^{d}$ | $3.6 \pm 0.4^{c}, (6.1 \pm 0.5)\times10^{3\ d}$ | $5.1 \pm 1.0^{d}, (6.4 \pm 0.4)\times10^{4\ c}$ | $5.2 \pm 2.8^{c}, (6.2 \pm 0.2)\times10^{5\ b}$ | $3.7 \pm 0.6^{c}, (5.7 \pm 0.1)\times10^{6\ a}$ |
| $1\times10^{2}$ | $61.1 \pm 5.9^{c}, 0^{d}$ | $52.9 \pm 2.5^{d}, 4.3 \pm 0.9^{d}$ | $45.6 \pm 3.8^{c}, 53.0 \pm 2.3^{d}$ | $48.1 \pm 2.2^{d}, 610.0 \pm 36.1^{d}$ | $57.3 \pm 4.7^{c}, (6.2 \pm 0.2)\times10^{3\ cd}$ | $48.8 \pm 4.2^{d}, (6.2 \pm 0.3)\times10^{4\ c}$ | $43.1 \pm 2.7^{c}, (6.0 \pm 0.1)\times10^{5\ b}$ | $47.0 \pm 6.0^{c}, (5.8 \pm 0.1)\times10^{6\ a}$ |
| $1\times10^{3}$ | $639.4 \pm 15.2^{c}, 0^{c}$ | $613.8 \pm 19.3^{d}, 4.1 \pm 1.1^{c}$ | $572.7 \pm 28.1^{c}, 45.5 \pm 4.1^{c}$ | $522.5 \pm 33.8^{d}, 562.7 \pm 11.2^{c}$ | $490.1 \pm 49.8^{c}, (6.0 \pm 0.3)\times10^{3\ c}$ | $518.3 \pm 17.0^{cd}, (6.2 \pm 0.4)\times10^{4\ c}$ | $489.1 \pm 18.7^{c}, (5.7 \pm 0.1)\times10^{5\ b}$ | $461.0 \pm 15.8^{c}, (5.8 \pm 0.1)\times10^{6\ a}$ |
| $1\times10^{4}$ | $(6.0 \pm 0.2)\times10^{3\ c}, 0^{d}$ | $(6.1 \pm 0.4)\times10^{3\ d}, 5.5 \pm 1.7^{d}$ | $(5.5 \pm 0.2)\times10^{3\ c}, 46.1 \pm 12.1^{d}$ | $(5.2 \pm 0.1)\times10^{3\ cd}, 591.1 \pm 71.5^{d}$ | $(5.5 \pm 0.2)\times10^{3\ c}, (6.0 \pm 0.6)\times10^{3\ d}$ | $(5.3 \pm 0.3)\times10^{3\ cd}, (6.0 \pm 0.2)\times10^{4\ c}$ | $(5.5 \pm 0.3)\times10^{3\ c}, (5.7 \pm 0.2)\times10^{5\ b}$ | $(5.2 \pm 0.2)\times10^{3\ c}, (5.5 \pm 0.1)\times10^{6\ a}$ |
| $1\times10^{5}$ | $(6.1 \pm 0.2)\times10^{4\ c}, 0^{d}$ | $(6.1 \pm 0.2)\times10^{4\ c}, 4.7 \pm 1.5^{d}$ | $(5.8 \pm 0.1)\times10^{4\ c}, 49.4 \pm 7.7^{d}$ | $(5.7 \pm 0.2)\times10^{4\ c}, 586.1 \pm 95.6^{d}$ | $(5.5 \pm 0.3)\times10^{4\ c}, (5.7 \pm 0.2)\times10^{3\ d}$ | $(5.4 \pm 0.3)\times10^{4\ cd}, (5.7 \pm 0.2)\times10^{4\ c}$ | $(5.4 \pm 0.4)\times10^{4\ c}, (5.3 \pm 0.2)\times10^{5\ b}$ | $(4.9 \pm 0.2)\times10^{4\ c}, (5.4 \pm 0.1)\times10^{6\ a}$ |
| $1\times10^{6}$ | $(6.3\pm0.1)\times10^{5\ b}, 0^{d}$ | $(6.1 \pm 0.1)\times10^{5\ b}, 5.2 \pm 1.2^{d}$ | $(6.0 \pm 0.1)\times10^{5\ b}, 50.7 \pm 8.2^{d}$ | $(5.7 \pm 0.1)\times10^{5\ b}, 544.9 \pm 60.2^{d}$ | $(5.6 \pm 0.1)\times10^{5\ b}, (5.4 \pm 0.3)\times10^{3\ d}$ | $(5.4 \pm 0.1)\times10^{5\ b}, (5.2 \pm 0.2)\times10^{4\ c}$ | $(5.2 \pm 0.1)\times10^{5\ b}, (5.2 \pm 0.1)\times10^{5\ b}$ | $(4.9 \pm 0.1)\times10^{5\ b}, (5.2 \pm 0.0)\times10^{6\ a}$ |
| $1\times10^{7}$ | $(6.4\pm0.1)\times10^{6\ a}, 0^{d}$ | $(6.2 \pm 0.1)\times10^{6\ a}, 4.4 \pm 01.6^{d}$ | $(5.7 \pm 0.1)\times10^{6\ a}, 46.8 \pm 5.5^{d}$ | $(4.7 \pm 0.1)\times10^{6\ a}, 513.5 \pm 55.0^{d}$ | $(4.8 \pm 0.1)\times10^{6\ a}, (5.1 \pm 0.4)\times10^{3\ cd}$ | $(4.8 \pm 0.1)\times10^{6\ a}, (5.1 \pm 0.2)\times10^{4\ c}$ | $(4.9 \pm 0.1)\times10^{6\ a}, (5.0 \pm 0.1)\times10^{5\ b}$ | $(4.9 \pm 0.1)\times10^{6\ a}, (4.8 \pm 0.1)\times10^{6\ a}$ |

Data are presented as mean ± SD, n = 3. abcd, data with different symbols are significantly different.

**Supplementary experimentals**

**1. Reagents, strains, and fecal samples**

All regular reagents were purchased from domestic merchandisers, Merck KGaA (Darmstadt, Germany) or ThermoFisher Scientific (Waltham, USA). The ssDNA libraries and aptamers were synthesized by Sangon Biotech Co., Ltd (Shanghai, China). The cloning pMD$^{TM}$19-T simple vector was purchased from Takara Co., Ltd. (Shiga, Japan). Furthermore, all bacterial strains were obtained from the American Type Culture Collection, including *Aliarcobacter butzerli* ATCC 49616, *Aliarcobacter skirrowii* ATCC 51132, *Aliarcobacter cryaerophila* ATCC 43158, *Campylobacter jejuni* ATCC 33560, *Campylobacter coli* ATCC 33559, *Campylobacter fetus* ATCC 27374, *Helicobacter pylori* ATCC 700392, *Lactobacillus casei* ATCC 393, *Lactobacillus johnsonii* ATCC 11506, *Bifidobacterium bifidum* ATCC 29521, *Enterococcus faecalis* ATCC 19433, *Escherichia coli* ATCC 11775, *Salmonella enteritidis* ATCC 25928, *Listeria monocytogenes* ATCC 19115, *Staphylococcus aureus* ATCC 12600, *Streptococcus gordonii* ATCC 35557, *Ralstonia pickettii* ATCC 27511 and *Shigella dysenteriae* ATCC 13313. The strains of the species belong to *Campylobacter* and *Aliarcobacter* genera were cultured on Columbia Blood Agar Base (Solarbio, China) containing 7% defibrinated horse blood for 48 h at 37°C under microaerophilic conditions. *H. pylori* were cultured on Brain Heart Infusion (BHI) Agar for 48 h at 37°C under microaerophilic conditions [4]. *L. casei*, *B. bifidum*, *L. johnsonii*, *E. faecalis* were cultured on MRS medium for 48 h at 37°C under anaerobic conditions. *E. coli*, *S. enteritidis*, *L. monocytogenes*, *S. gordonii*, *R. Pickettii*, *S. aureus* and *S. dysenteriae* strains were cultured in Luria-Bertani medium under aerobic conditions. The cuitivation and operations of bacteria categorized as biosafty level 2 were implemented in the P2 laboratory according to the regulations. Additionally, fecal samples were provided by Qingdao Municipal

Hospital from *A. butzerli*-, *C. jejuni*- or *C. coli*-uninfected children. The chicken broiler was purchased from a supermarket.

### 2. Preparation and characterization of fusarinine C[$Fe^{3+}$] chelates-modified magnetic nanoparticles

$Fe^{3+}$-chelating fusarinine C (FsC[$Fe^{3+}$]) was obtained following the methods described previously [5]. MNPs was prepared with formerly described protocols [6]. To modify MNPs with FsC[$Fe^{3+}$], 1.5 mg FsC[$Fe^{3+}$] was reacted with 1 mg MNPs via the catalysis of 3.8 mg 1-Ethyl-3-(3-dimethylaminopropyl) carbodiimide hydrochloride (EDC·HCl) and 4.6 mg N-hydroxysuccinimide (NHS) in 100 mL 15 mM MES buffer (pH 6) for 48 h at 4℃. The reacted MNPs were collected by magnetic separation, and 1 mg of them were washed three times with de-ionized $H_2O$ and solidified with freeze drying. The characteristic bonds of the obtained black powder were analyzed by FT-IR; the surface charge was measured by a Malvern ZEN3600 Zetasizer Nano instrument (Malvern, UK); its morphology and average particle size was determined by Transmission Electron Microscopy (TEM) and Nano Measurer software [6]. To block the free surface carboxyl groups on the FsC[$Fe^{3+}$]-MNPs, 1 mg of them were incubated with 5.0% $CaCl_2$ [6] in 1 mL 1×PBS buffer for 60.0 min and washed twice with 1× PBS buffer. After freeze drying, the $Ca^{2+}$-doped FsC[$Fe^{3+}$]-MNPs (Ca/FsC[$Fe^{3+}$]-MNPs) was obtained.

### 3. Selectivity of Ca/FsC[$Fe^{3+}$]-MNPs to bacteria

1 mg Ca/FsC[$Fe^{3+}$]-MNPs were resuspend in 100 mL of 15 mM MES buffer (pH 6.0). Then, ultrasonic homogenization for 5 min was performed to fully disperse them, obtaining a transparent solution. The specificity of Ca/FsC[$Fe^{3+}$]-MNPs for *C. jejuni* and *A. butzleri* were determined by respectively incubating Ca/FsC[$Fe^{3+}$]-MNPs with artificial bacterial communities, which were the one

contained *C. jejuni*, *A. butzleri* and some Gram-negative bacteria and the other of these two bacteria mixed with other Gram-positive bacteria. In detail, 1 mg Ca/FsC[$Fe^{3+}$]-MNPs was incubated with a bacterial community containing *C. jejuni*, *A. butzleri*, *E. coli*, *S. enteritidis*, *R. pickettii* and *S. dysenteriae* ($10^7$ CFU/mL each) for 30 min at 37°C in 1mL 1× PBS buffer, and then collected by magnetic separation. The magnetically separated products was transferred to 200.0 μL de-ionized water, and then incubated in boiled water bath for 30 min; Meanwhile, 1 mg Ca/FsC[$Fe^{3+}$]-MNPs was incubated with the community containing *C. jejuni*, *A. butzleri* and other 6 gram-positive bacteria of *L. casei*, *B. bifidum*, *L. johnsonii*, *E. faecalis*, *L. monocytogenes* and *S. gordonii*, and the magnetically separated products were treated with 2 mg/mL lysozyme in 200.0 μL 1mL 1× PBS buffer at 37°C for 30 min to hydrolyze the cell walls of gram-positive bacteria. 16S rDNA PCR amplification [7] was implemented to the treated supernatant of the collected products, respectively. The resulting correct fragment from each PCR product was recombined into the pMD$^{TM}$19-T simple vector, transformed into *E. coli* DH5α, and 200 clones were picked out for Sanger sequencing by Sangon Biotech (Shanghai, China). The resulting sequences were aligned by nucleotide BLAST at NCBI website to identify the specific species of the 200 clones, and the number of each species was counted.

**4. Preparation and characterization of aptamer-modified CDs**

Two different colored carbon dots were selected to modify the aptamers in this dual-bacteria detection biosensor module. Red emitting carbon dots (RCDs) [1] and green emitting carbon dots (GCDs) [2] were prepared using the method as previously reported. The two CDs were synthesized to verify their luminescence performance by UV-Vis absorption spectroscopy and fluorescence spectroscopy. Moreover, their surface functional groups were characterized by FT-IR. Subsequently, the shape and size of the particles are observed by TEM.

For the aptamer-modified CDs, 5'-amine modified aptamers were synthesized by Sangon Biotech (Shanghai, China), and they were conjugated to the carboxyl groups of their corresponding CDs catalyzed by EDC·HCl/NHS. In specific, 200 μg CDs were incubated with 4.8 mg EDC·HCl and 2.9 mg NHS in 1 mL 1×PBS buffer (pH7) at 25°C for 1 h. Afterwards, 5' amine-aptamer and carboxyl-activated CDs were incubated at a series of mole/weight ratios in 1mL 1×PBS buffer at 25°C for 2 h, which were 20, 40, 60, 80, 100 and 120 pmol/μg for *C. jejuni*-specific aptamer and RCDs, and 15, 30, 45, 60, 75 and 90 pmol/μg for *A. butzleri*-specific aptamer and GCDs; then, they were transferred to 4°C and kept for another 12 h. Afterwards, they were centrifuged using an ultrafiltration centrifuge tube (Mw cut-off = 20,000) to separate un-reacted aptamers, and they were quantified by using SYBR Green II dye [8]. The weight of grafted aptamers was determined by subtracting the unreacted amounts from the initial ones, thus to calculate the grafting ratios of aptamers on CDs. The aptamer-modified CDs were freeze dried and subjected to the characterizations by FT-IR and electrophoretic mobility shift assay (EMSA) [9]. To validate the specificity of *C. jejuni*-aptamer-CDs for this bacterium, 7.5 μg of the CDs was incubated with *C. jejuni* and other 6 control bacteria of *E. coli*, *R. pickettii*, *S. enteritidis*, *L. johnsonii*, *E. faecalis* and *B. bifidum* (each at the density of $1.0 \times 10^7$ CFU/mL) at 37°C for 15 min, and then tested for fluorescence as described above. Similarly, the specificity of *A. butzleri*-aptamer-CDs was validated with the same procedures only by changing the bacterium to *A. butzleri*.
